# *Ribo-ODDR*: Oligo Design pipeline for experiment-specific Depletion of Ribosomal RNAs in Ribo-seq

**DOI:** 10.1101/2020.01.12.900175

**Authors:** Ferhat Alkan, Joana Silva, Eric Pintó Barberà, William J. Faller

## Abstract

Ribosome profiling (Ribo-seq) has revolutionized the study of RNA translation by providing information on ribosome positions across all translated RNAs with nucleotide-resolution. Yet, several technical limitations restrict the sequencing depth of such experiments, the most common of which is the overabundance of ribosomal RNA (rRNA) fragments, which frequently make up more than 90% of sequencing reads if not depleted. Various strategies can be employed to tackle this issue, including the use of commercial rRNA depletion kits. However, as they are designed for more standardized RNAseq experiments, such kits may perform suboptimally in Ribo-seq. There is therefore potential to significantly increase the information that can be gleaned from Ribo-seq experiments. Here we show that a major confounding issue is that the rRNA fragments generated via Ribo-seq vary significantly with differing experimental conditions, suggesting that a “one-size-fits-all” approach may result in inefficient rRNA depletion. In order to overcome this, it is possible to use custom-designed biotinylated oligos complementary to the most abundant rRNA fragments, however currently no computational framework exists to aid the design of optimal oligos. We have developed *Ribo-ODDR*, an oligo design pipeline integrated with a user-friendly interface that assists in oligo selection for efficient experiment-specific rRNA depletion. *Ribo-ODDR* uses preliminary data to identify the most abundant rRNA fragments, and calculates the rRNA depletion efficiency of potential oligos. We show that *Ribo-ODDR* designed oligos lead to a significant increase in rRNA depletion, and increased sequencing depth as a result, providing substantial information that would otherwise have been lost. *Ribo-ODDR* is freely accessible at https://github.com/fallerlab/Ribo-ODDR

## 1 Introduction

Since its development, Ribosome Profiling (also known as Ribo-seq) has revolutionized the study of RNA translation [1]. The technique allows the analysis of ribosomally associated mRNA at codon-level resolution, providing a snapshot of the mRNAs bound by ribosomes in the cell. Information on translation efficiencies, open reading frame (ORF) usage, translation start sites, ribosome pause sites, amino acid dependencies, and translation elongation rates can be gleaned from the data generated (reviewed in [2]). Additionally, the level of ribosome binding to an mRNA is a much better predictor of protein levels than the quantity of mRNA that is present, underscoring the importance of this technique [1, 3].

The Ribo-seq protocol takes advantage of the fact that at any instant a ribosome covers a *∼*28 nucleotide fragment of mRNA. This fragment is protected from nuclease digestion as a result and is hence known as the ribosome protected fragment (RPF). Following ribosome stalling with translation blockers (i. e. cycloheximide), isolation of a cell lysate, and treatment with RNase, a cDNA library can be made from the resulting RPFs, and sequenced. By selecting the correct fragment size, the abundance of ribosomes at every location on the transcriptome can be deduced.

Although this process has been somewhat standardized [4], it is acknowledged that numerous problems remain in generating high quality data. The RNase enzyme used [5], or length of digestion [6] can significantly bias the resulting data. Additionally, it is a common problem that a high proportion of sequencing reads derive from rRNA sequences, despite the use of rRNA depletion strategies. Indeed, in most experiments rRNA make up the majority of all reads sequenced [4], and more than 90% in some cases [7].

At present, the most common rRNA depletion strategies include the use of commercial rRNA depletion kits or the use of custom-designed biotinylated oligos previously reported in the literature. Both of these approaches make use of RNA oligos that are complementary or near-complementary to the rRNA, thus binding to their target rRNA sequence and allowing its depletion with a simple fishing approach. Additionally, the use of duplex-specific nuclease (DSN) has been reported [8]. However, DSN is known to also deplete highly expressed genes, and both commercial kits and custom oligos assume that the rRNA fragments present in a sample are consistent across experiments. Here we show that this is not the case, and that the experimental conditions and the tissue being used both introduce variations in the abundance of rRNA fragments produced. This raises the possibility that differential efficiencies of rRNA depletion across samples in an experiment may introduce biases in Ribo-seq data [9].

There are a number of possible approaches that could be taken to circumvent this problem. For example, there may be previously published data that provides a list of oligos that is confirmed to be efficient for Ribo-seq performed in a specific tissue and organism, following a specific protocol. However these pre-designed oligos may need further optimization. For example, the overall efficiency of pre-designed oligos can be improved by their cross-species optimization. The most reliable way to confront the problem of differential rRNA fragmentation is to perform pilot experiments on identical or similar samples and design novel biotinylated oligos that targets the most abundant rRNA fragments within generated pilot data [10, 9]. Unfortunately, this approach requires experimental effort and computational work, potentially with a few rounds of optimization. However, this could be avoided due to the increasing number of Ribo-seq datasets from diverse sources that are being published, which could also serve as pilot data for researchers. Using this data, the most abundant rRNA fragments can be identified, and oligos designed to deplete them.

With this study, we first provide evidence that commercial rRNA depletion kits perform suboptimally and rRNA fragments generated by nuclease treatment differ substantially under various experimental conditions. Furthermore, we show that the same variability exists in fragments generated from different organs, even when using identical protocols. To tackle this problem, we present *Ribo-ODDR*, a Ribo-seq focused Oligo Design pipeline for Depleting rRNAs. This pipeline addresses and automates the above mentioned problems and allows the design or optimization of oligos with high rRNA depleting potential, based on preliminary or previously published data. It is freely accessible via GitHub in order to help researchers improve the power of their Ribo-seq experiments through more efficient rRNA depletion, thus maximizing the information gained from Ribo-seq experiments.

## 2 RESULTS

### 2.1 Suboptimal rRNA depletion of commercial rRNA depletion kits

Inefficient rRNA depletion is a known issue in Ribo-seq, and recently a comparative analysis of different rRNA depletion approaches has been published [9]. This analysis included several commercially available kits (Ribo-Zero, Ribo-Zero Plus, RiboCop, NEBNext, and QIAseq FastSelect), as well as a pool of biotinylated custom oligos (riboPool). Surprisingly, analysis of this data showed that despite rRNA depletion, there was still a high abundance of rRNA fragments in all samples. Of all reads that could be mapped to rRNA and protein coding transcripts, an average of 85% of them were rRNA fragments (see Figure 1). These unexpectedly high percentages significantly reduce the resolution of the performed experiments as they decrease the sequencing depth in open reading frames (ORFs) and thus limit downstream analyses.

**Figure 1:**
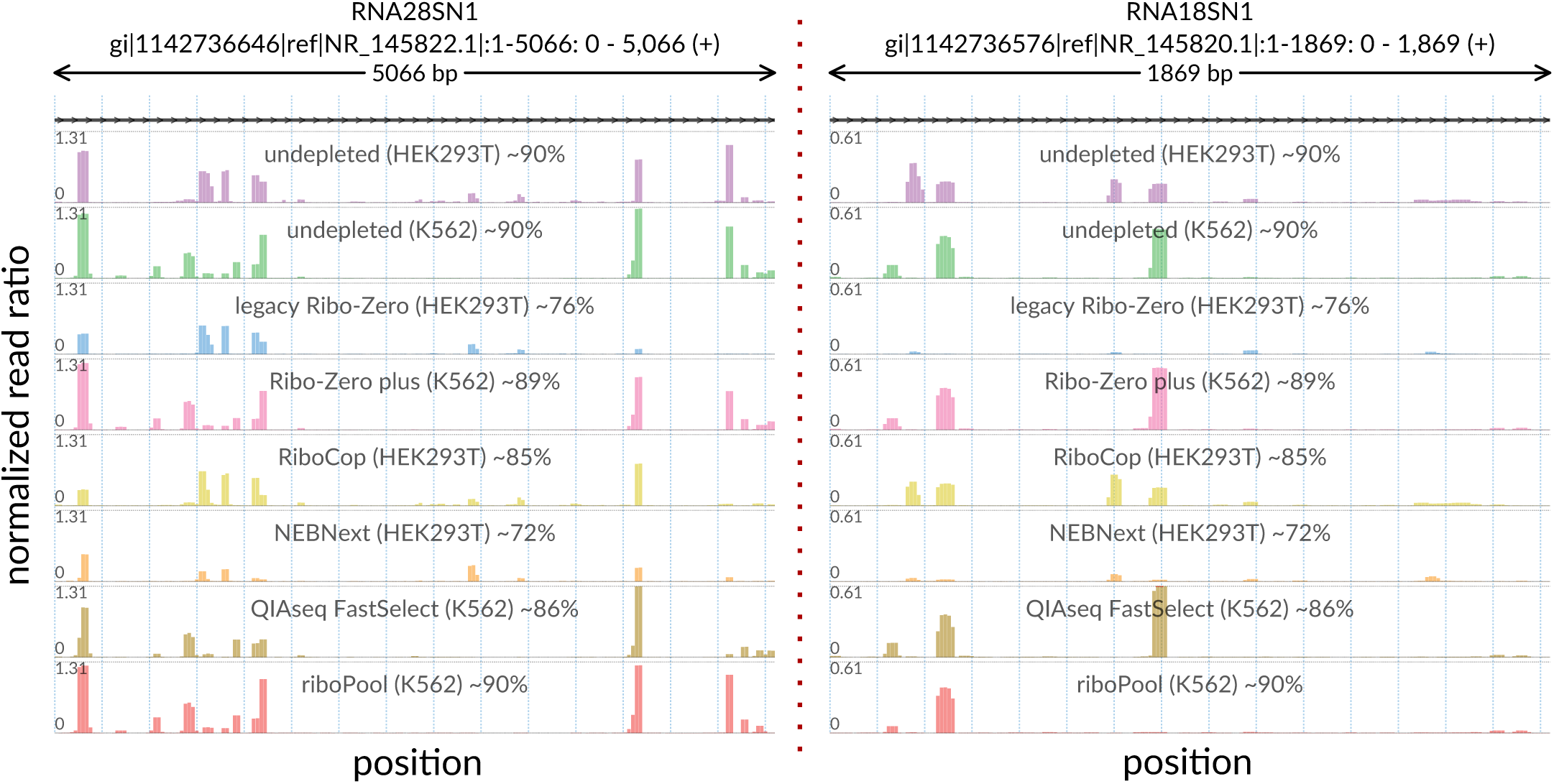
Suboptimal performance of commercial rRNA depletion kits. Visualization is based on a previously published dataset where several different kits were tested for rRNA depletion in human cell lines. Each track shows the positional abundance profile of 28S (left) and 18S (right) rRNA fragments coming from individual samples. For every position in the x-axis, y-axis represents the normalized read ratio, number of rRNA reads mapped to that position divided by the total number of reads mapped to all protein coding transcripts. Sample-specific total rRNA percentages are given in track labels together with sample identifiers.

In order to further understand the inefficiencies of these kits, we visualized this data using the *svist4get* tool [11]. In line with the published analysis, we observed that rRNA depletion using commercial kits resulted in the incomplete depletion of 28S and 18S rRNA fragments, particularly those originating from several experiment-specific hotspots within each. Interestingly however, there was significant variability in the rRNA fragments present, suggesting that there was a depletion protocol driven heterogeneity in the rRNA fragments sequenced (Figure 1). Analysis of an additional dataset generated using Ribo-Zero [12] also showed this heterogeneity, suggesting experiment-specific rRNA inconsistencies introduce variables that result in decreased protein coding sequencing depth (see Supplementary Figure 1).

In samples that are difficult to work with, this may be a terminal issue for profiling experiments. Ribo-seq in intestinal epithelial cells, for example, is known to result in a very high level of rRNA sequencing reads (unpublished data from this lab and personal communication from others). We therefore carried out a Ribo-seq experiment using *in vitro* mouse intestinal organoids, with rRNA fragments depleted with the RiboCop kit. This experiment resulted in 89% rRNA reads (only 11% protein coding reads). In our analysis, we identified three hotspots (one for each 28S, 18S and 5-8S rRNA), where each hotspot individually accounted for more sequencing reads than all protein-coding transcript reads (see Supplementary Figure 2).

**Figure 2:**
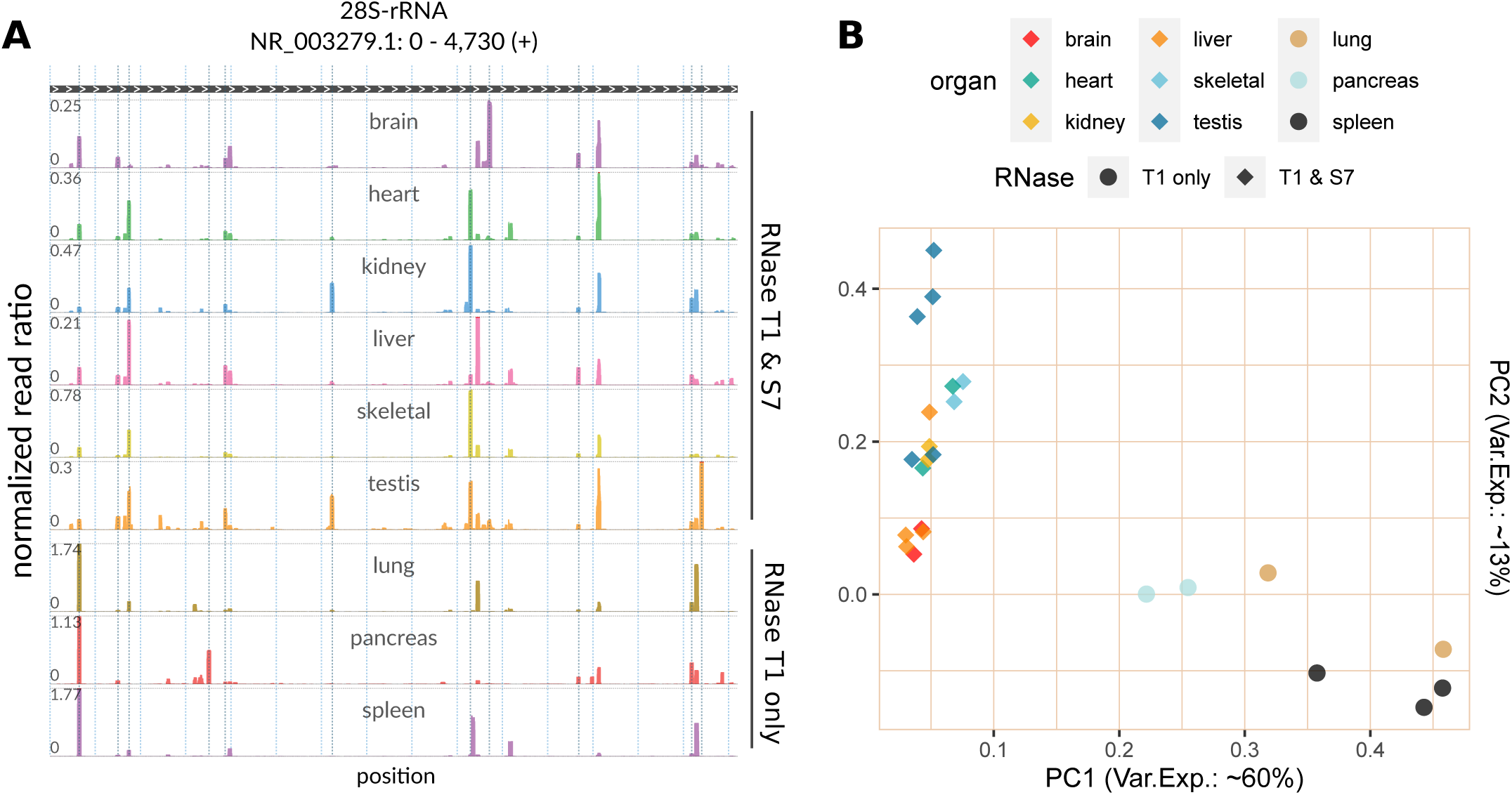
Tissue and RNase specificity of rRNA fragments in the Ribo-seq data from [13]. (A) Each track shows the positional abundance profile of 28S rRNA fragments within the representative sample of the labeled organ. For every position in the x-axis, y-axis represents the normalized read ratio, number of rRNA reads mapped to that position divided by the total number of reads mapped to all protein coding transcripts. (B) Using the sample-specific abundance profiles on all rRNAs (28S, 18S, 5-8S and 5S), principal component analysis (PCA) was performed for all samples of the analyzed dataset (number of replicates varies for each organ). The first and second principal components are plotted against each other, PC1 in x-axis and PC2 in y-axis. The percentage of variance explained by each component is given in corresponding axis labels.

These observations demonstrate that commercial depletion kits perform suboptimally in Ribo-seq and suggest that custom-designed rRNA depletion oligos would be a powerful way to increase sequencing depth in such experiments.

### 2.2 Tissue and RNase specificity of rRNA fragments in mouse

The use of custom-designed biotinylated oligos serves as a good alternative to overcome the inefficiency of commercial rRNA depletion kits in Ribo-seq experiments. However, there is no consensus on which oligos to use for maximal rRNA depletion, or even whether the same oligos are suitable for different experiments. Our results above would suggest that this is not the case.

In order to assess this, we measured the variability in rRNA fragment position and abundance in samples generated using different protocols and tissues of origin. We made use of a previously published dataset in which the authors performed *in vivo* Ribo-seq in nine different mouse organs without any rRNA depletion [13]. In this dataset, six sets of samples (brain, heart, kidney, liver, skeletal muscle, and testis) were digested using a mix of RNaseT1 and RNaseS7, with the remaining 3 (lung, pancreas, and spleen) being digested with only RNaseT1 as part of the Ribo-seq protocol. After identifying 28S, 18S, 5-8S and 5S rRNA fragments separately for each sample, we compared their rRNA fragment profiles (based on number of fragments mapped to each position in rRNAs) with a principal component analysis (PCA) (see Figure 2). This analysis revealed a striking heterogeneity in rRNA fragments in samples generated using different protocols, suggesting that rRNA depletion oligos that are efficient in one experiment may not be suitable for another. This protocol-derived heterogeneity of rRNA fragments can be clearly observed in Figure 2A, where the positional abundance profiles 28S rRNA fragments are shown for individual organs (one representative sample for each).

Moreover, the PCA and abundance profiles also reveal significant rRNA fragment differences in samples generated from different organs even when using the same protocol. While our analyses showed that there is a strong agreement between replicate measures of each organ in terms of rRNA fragments produced (Supplementary Figure 3-11), we observed clear profile separation between organs (Figure 2 and Supplementary Figure 12). This suggests that rRNA fragment heterogeneity is a common occurrence, and clearly shows that a “one size fits all” approach is not appropriate for rRNA depletion in Ribo-seq experiments.

**Figure 3:**
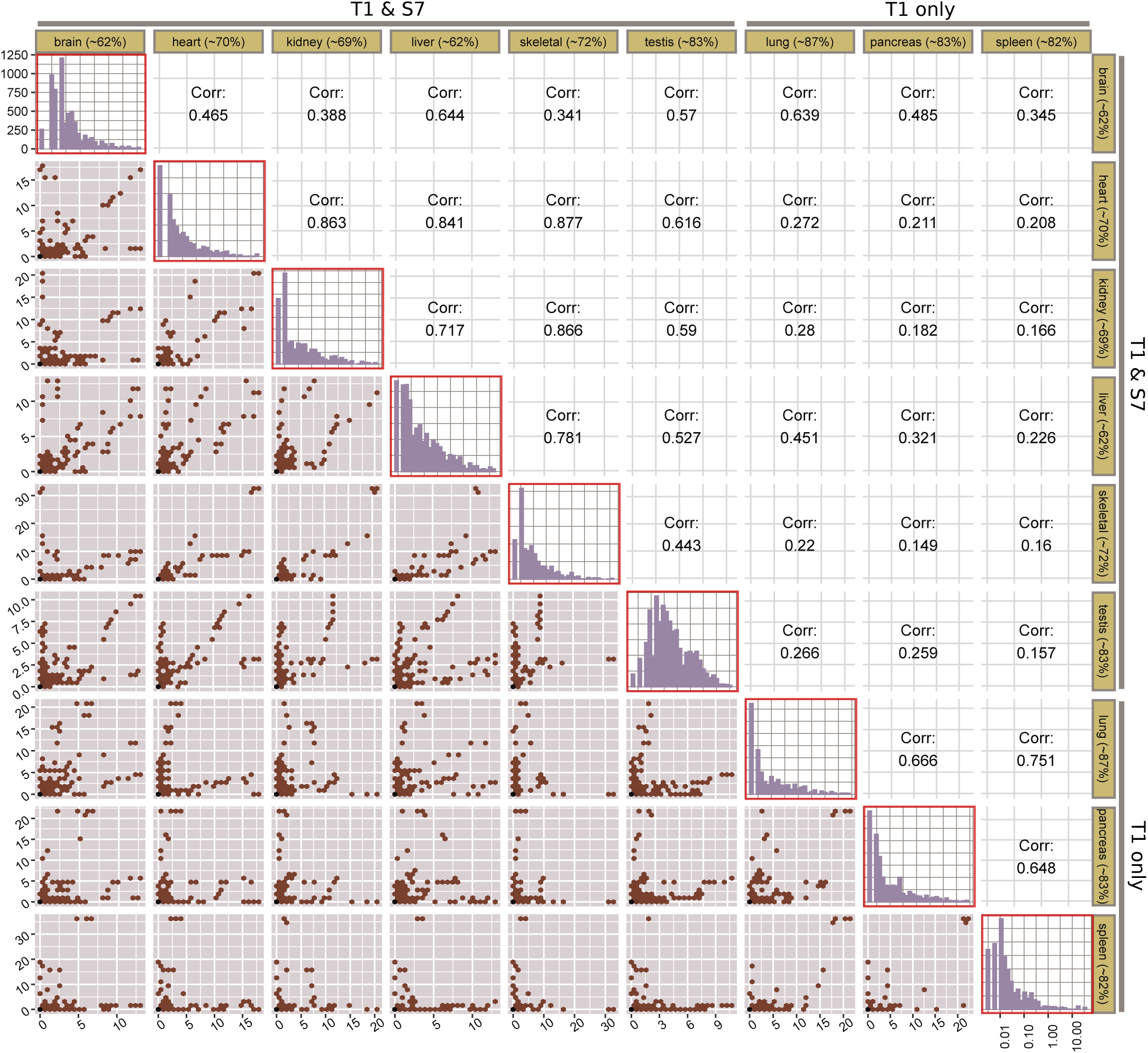
Cross-organ correlation analysis of *Ribo-ODDR* computed oligo depleting potentials for all 25nt oligos (n=6782) targeting mouse 28S, 18S, 5-8S and 5S rRNA fragments. Each row and column corresponds to an organ. The diagonal plots (red boxes) show the histogram of oligo depleting potentials, computed for that organ based on the analyzed dataset. Axes of diagonal plots are shared, x-axis (shown in bottom-right corner) representing the log-transformed depleting potential and y-axis (shown in top-left corner) representing the number of oligos with that potential. Lower hex-binned scatter plots compares the depleting potential of all oligos between organ pairs (column vs row) with the Pearson’s correlation coefficient given in their diagonal mirrors. In these plots, each bin contains one or more oligos with organ-specific rRNA depleting potential given in x- and y-axes for column and row organs, respectively. Percentages in row and column labels show the average rRNA percentage for that organ.

### 2.3 Comparing the depleting potential of oligos across different experiments

In order to understand the effect that this rRNA fragment heterogeneity has on the efficiency of rRNA depletion oligos, we developed *Ribo-ODDR*. Based on given pilot Ribo-seq data, this pipeline measures the depleting potential of all possible oligos. For each oligo, this potential is simply equal to the percentage of rRNA fragments produced from the oligo target region on the rRNA, where the oligo sequence binds with near-perfect complementarity (see Materials and Methods).

We ran *Ribo-ODDR* on the organ-specific data used above and obtained the sample-specific depleting potentials of all 25nt long oligos (*n* =6782) that can deplete mouse rRNA fragments. For each individual oligo, the organ-specific depleting potential was calculated by simply averaging the values computed for each replicate of that organ.

In Figure 3, we compare the depleting potentials of oligos across all organ pairs with a cross-organ correlation analysis. These data makes it clear that the correlation in oligo depleting potential between samples treated using the same RNase digestion strategy is significantly higher than those using another strategy. In the RNaseT1-only digestion group, intra-group Pearson’s correlation coefficients are between 0.64 and 0.76 (mean value of 0.69), and for the RNaseT1/S7 group this is between 0.34 and 0.88 (mean value of 0.64). This confirms our observations detailed in Figure 2 that differing experimental conditions results in substantial differences in rRNA fragments created.

Furthermore, if the same RNase digestion protocol is used, oligos designed for one tissue (assuming only high potential oligos are selected) do not necessarily provide efficient depletion in another. In some cases, the oligos with high depleting potential in one tissue show high depleting potential in others (kidney-*vs*-skeletal muscle, for example), however, this is only the case for a minority of tissues. Most tissue pairs show a low correlation in rRNA depleting potential of oligos. For example, pancreas-*vs*-heart and skeletal muscle-*vs*-lung both have Pearson’s correlation coefficients of below 0.25, demonstrating that rRNA depletion oligos used successfully in one organ are unlikely to work in another. These observations are in agreement with our previous analysis of other publically available datasets (Figure 1), and suggest that maximizing the information gained in Ribo-seq experiments may require experiment-to-experiment optimization.

### 2.4 Improving overall rRNA depletion efficiency using *Ribo-ODDR, in vivo* oligo design example

To test the power of the *Ribo-ODDR* design platform, we performed *in vivo* Ribo-seq experiments in the mouse intestine (see Figure 4).

**Figure 4:**
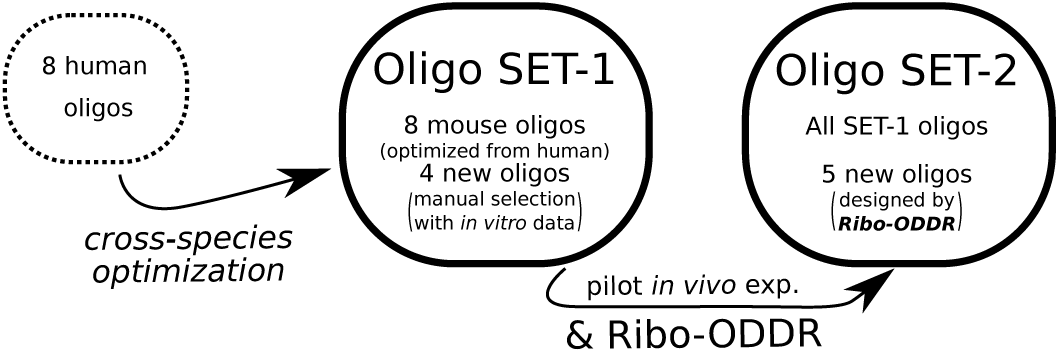
Oligo sets used in this paper. SET-1 consists of mouse-optimized version of eight human oligos and four additional ones, manually selected based on pilot *in vitro* experiments. SET-2 includes all SET-1 oligos and 5 new oligos, designed by *Ribo-ODDR* with data from pilot *in vivo* experiments using SET-1 oligos for rRNA depletion.

We began by optimizing previously published human oligos [1, 14] to mouse ribosomal sequences which we named SET-1 oligos (see Supplementary Figure 13 and Supplementary Table 1-2). This experiment resulted in an average of only *∼*6% of sequencing reads mapped to protein coding regions, confirming the high levels of rRNA contamination found in intestinal epithelial samples. With this pilot data, we ran the *Ribo-ODDR* pipeline to design 5 additional oligos with high rRNA depleting potentials and added them to the oligo pool, creating SET-2 (see Supplementary Table 3). In Figure 5 and Supplementary Figure 14, we show that positional abundance profile of rRNA fragments are highly conserved between replicates in each experiment group, and newly designed oligos in SET-2 were successful at depleting the fragments in their corresponding regions. Crucially, rRNA depletion was far more efficient after the addition of five *Ribo-ODDR* designed oligos, resulting in a *∼*5-fold increase in protein-coding transcript reads (*∼*28% vs *∼*6%), with SET-2 oligos giving *∼*72% rRNA fragments on average, compared to *∼*94% rRNA fragments on average for experiments using SET-1 oligos. This substantial increase in rRNA depletion efficiency demonstrates the power of experiment-specific rRNA depletion in Ribo-seq experiments and how using *Ribo-ODDR* can help this process.

**Figure 5:**
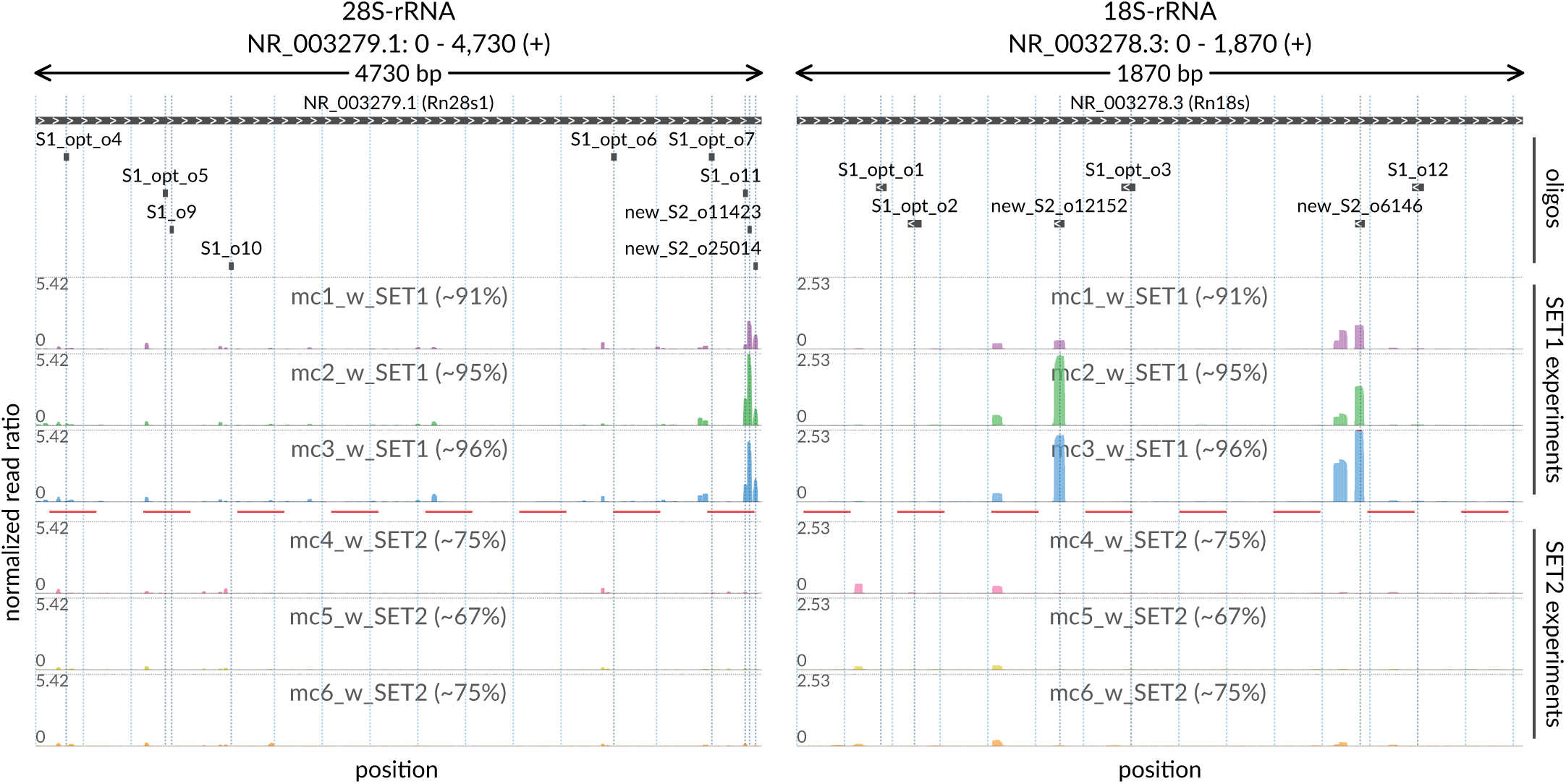
Positional abundance profiles of 28S (left) and 18S (right) rRNA fragments coming from *in vivo* (mouse intestine) Ribo-seq experiments performed with two different sets of rRNA depletion oligos, SET-1 and SET-2. The latter set includes all oligos from the former and 5 additional oligos designed with *Ribo-ODDR*, based on pilot data generated using only SET-1 oligos. In each figure, top track indicates the target regions of used oligos within that rRNA, where additional oligos of the SET-2 are labeled as ‘new’. In all tracks, x-axis corresponds to position within rRNAs. In profile tracks, y-axis is fixed for all samples and shows the normalized read ratio, number of rRNA reads mapped to the position divided by the total number of reads mapped to all protein coding transcripts. The percentages given within sample labels indicates the sample-specific percentage of rRNA fragments, within all reads that is mapped to rRNAs and protein-coding transcripts.

#### 2.4.1 Evaluating potential off-target effects of custom oligos

A potential drawback of oligo-based depletion of rRNA is the possibility of complementarity to protein-coding fragments that can result in off-target depletion of mRNA. To ensure that this is not the case with *Ribo-ODDR* designed oligos, the tool reports the off-target potential of all oligos, allowing the selection of those with minimal complementarity to mRNAs. Indeed, in-depth read count analysis of potential off-target sites, shows that such depletion can be avoided. As shown in Supplementary Figure 15, the average read count of potential off-target regions of the *Ribo-ODDR* designed SET-2 oligos does not change between experiments that use SET-1 or SET-2 oligos. This observation suggests that these 5 oligos, designed and selected with *Ribo-ODDR*, do not cause undesired depletion of informative off-target fragments.

### 2.5 *Ribo-ODDR* oligo-based depletion vs commercial kits

For comprehensive evaluation of rRNA depletion performances of commercial kits compared to *Ribo-ODDR* designed oligos, we performed six more *in vivo* Ribo-seq experiments in the mouse intestine, with 5 additional *Ribo-ODDR*-designed oligos, creating the SET-3. In the resulting data we measured the percentage of reads that map to the protein coding, ribosomal, intronic and other RNAs, as well as the size of the RNA fragments produced.

Analysis of these Ribo-seq experiments showed that RiboCop and RiboZero kits produced less than 5% protein coding mapped reads on average (see Figure 6), severely limiting the sequencing resolution of this experiment. However, *Ribo-ODDR* oligos yielded *∼*14% protein-coding mapped reads, showing *∼*3 to *∼*4 times better performance than commercial kits. Further analysis of these data showed that the commercial kits differed in the quality of sequencing reads, with the RiboCop kit resulting in a higher number of short sequencing reads compared to RiboZero or *Ribo-ODDR* designed oligos. These experiments clearly show that experiment-specific custom oligos are superior to commercially available kits for rRNA depletion in Ribo-seq experiments, and that *Ribo-ODDR* provides a suitable tool for the design of such oligos. Furthermore, we have shown that in samples that have a low sequencing depth of protein coding RNAs, this increased rRNA depletion can turn a failed experiment into a successful one.

**Figure 6:**
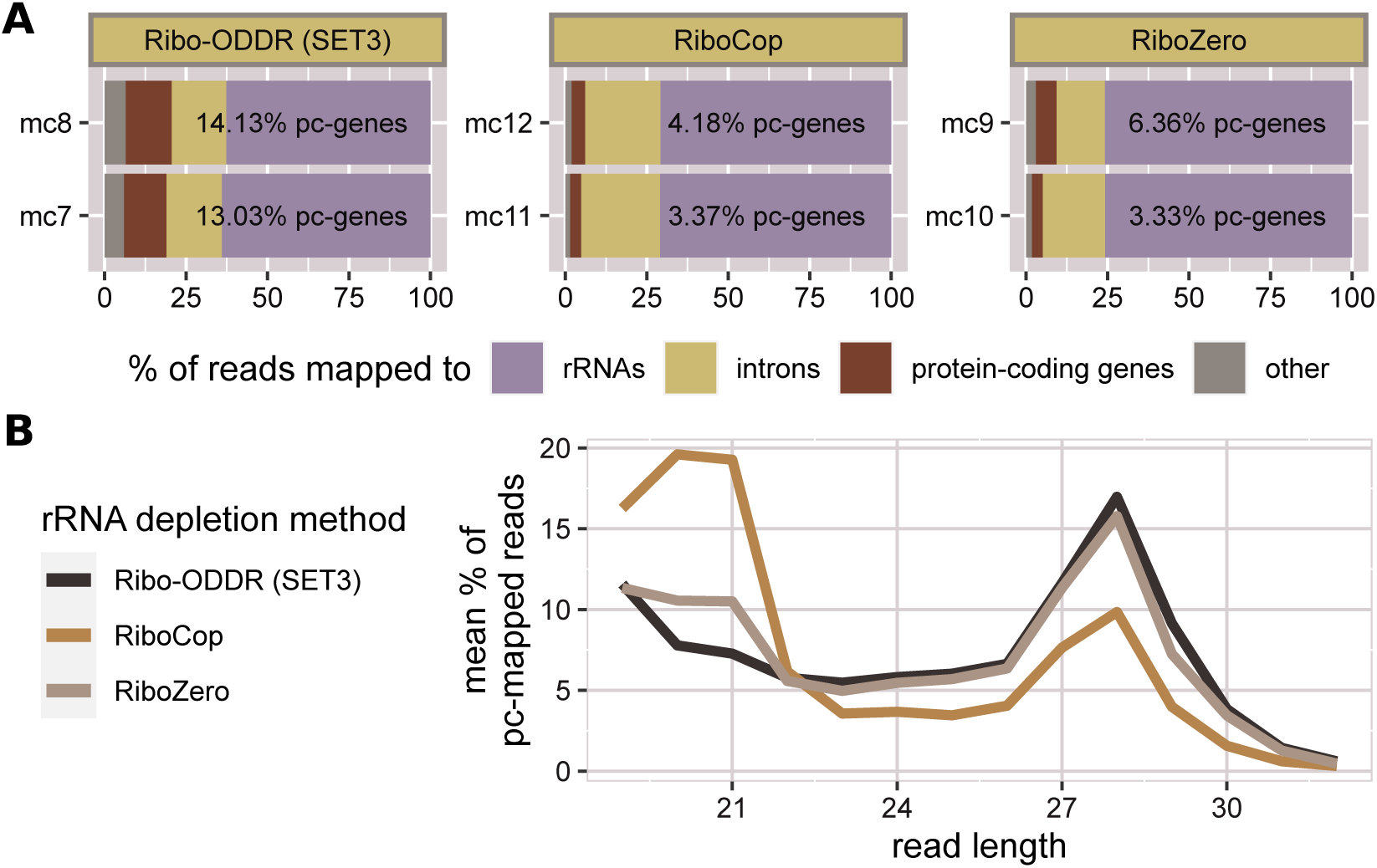
Performance comparison of *Ribo-ODDR* oligos (SET-3) with RiboZero and RiboCop rRNA depletion kits. (A) Fractions of sequencing reads that are mapped to protein-coding genes, rRNAs, other non-coding transcripts and introns in 6 Ribo-seq experiments, performed *in vivo* in mouse intestine (2 replicates per group). (B) Read length histogram for protein-coding gene mapped reads in 3 experiment groups. Percentages are given as average of two replicates for each group.

## 3 DISCUSSION

Ribosome profiling has become a mainstay experiment in the analysis of RNA translation. It is one of the most informative techniques available for studying the translatome and has become very widely used in the decade since its development [1]. However, as the technique focuses on ribosomally bound RNAs, the enrichment of rRNAs is an unfortunate necessity of the protocol. The nuclease cleavage of rRNAs produces fragments of a similar size to those being analyzed, creating an obvious technical challenge. Indeed, rRNA fragments commonly far outnumber reads from protein coding genes. As a result, rRNA depletion is a vital step in generating high quality Ribo-seq data.

The most common approach to overcome this issue is the use of commercially available rRNA depletion kits. However, our data shows that the efficiency of depletion using this method is variable, and suggests that combining this method with a small number of custom designed oligos could significantly increase rRNA depletion. Additionally, previously published studies have suggested that the use of commercial kits can result in bias in individual mRNA fragments [8, 9], emphasizing that the rRNA depletion strategy must be considered when planning experiments.

Using publically available data, we have also shown that this issue is compounded by variability in the specific rRNA fragments that is introduced by differing experimental conditions. Both the origin of the tissue, and the nuclease used for RNA digestion significantly change the rRNA fragment population, showing that a depletion strategy that works for one experiment will not necessarily work for another. The source of the tissue-specific rRNA fragment heterogeneity is unknown, however it may be due to differences in the accessibility to the tissue of the nuclease and/or to the presence of different intrinsic RNases. Ultimately, significant sequencing depth can be gained by improving the rRNA depletion. This may be particularly important in samples and tissues that have previously proven difficult to assay using Ribo-seq, such as the intestinal epithelium and other *in vivo* tissues.

We developed *Ribo-ODDR* to aid with the design of custom oligos in an experiment by experiment manner. The tool enables users to run the *design mode* using preliminary or previously published data, allowing them to select a number of high depletion potential oligos. We have shown that using such an approach can result in a 5-fold increase in the percentage of protein coding transcripts detected.

An obvious drawback of this approach is the need for preliminary data to optimize the depletion strategy. In order to optimally carry out a Ribo-seq experiment, it is advisable to generate such preliminary data using the exact protocol as planned under experimental conditions, particularly when using tissues that have previously proven difficult to work with. However, as a result of the increasing number of Ribo-seq studies being published, in many cases it may be sufficient to use data from a similar source tissue that has been previously published. This could then be analysed using *Ribo-ODDR* to create an oligo set that is likely to efficiently deplete rRNA.

Alternative depletion strategies have also been suggested, such as the use of duplex-specific nuclease (DSN) [8], which we have not compared to *Ribo-ODDR*-based depletion. However, it is important to point out that *Ribo-ODDR* is not necessarily a stand-alone method. We envision that *Ribo-ODDR* will be used alone in some cases, and in conjunction with other depletion strategies in others. For instance, our data suggests that commercial kits can benefit from the addition of a small number of custom designed oligos.

*Ribo-ODDR* gives experimenters a platform to assess the most optimal custom oligos, allowing for increased depth of mRNA fragment sequencing, and maximizing the information gained in Ribo-seq experiments.

## 4 MATERIALS AND METHODS

We first introduce the public datasets that are actively analyzed in this study to show the suboptimal performance of commercial rRNA depletion kits and rRNA fragment differences within different Ribo-seq experiments performed in the same species. Our analyses with these datasets serve as a justification for the experiment-specific depletion of rRNAs with custom oligos. Then, we describe the details of the *Ribo-ODDR* oligo-design pipeline and present its different modes of action. Finally, we explain the followed Ribo-seq protocol for the experiments performed within this study.

### 4.1 Ribo-seq with commercial rRNA depletion kits

Suboptimal performance of commercial rRNA depletion kits in Ribo-seq experiments is a known issue [8] and we provide evidence on this by analysing two public datasets [9, 12], in addition to one experiment performed here using the RiboCop kit (Lexogen, catalog no. 037). We accessed the public datasets through NCBI using GSE147324 (SRP253534) [9] and GSE96998 (SRP102438) [12] accession IDs and obtained the raw fastq files using the SRA Toolkit. For the former dataset, adapter trimming was performed following the instructions on the corresponding paper [9], and, for the latter, we used the cutadapt tool [15] for both adapter trimming and size selection marker cleaning. The rRNA fragments were then identified through mapping the trimmed read files to human and mouse 28S, 18S, 5.8S and 5S rRNA sequences, using the TopHat aligner [16]. We also mapped them to protein-coding transcript sequences (gencode release v34 and vM21) after cleaning rRNA fragments using the *SortMeRNA* tool [17], and calculated the rRNA percentages in the samples by dividing the number of rRNA-mapping reads by the total number of reads that maps to rRNAs or protein coding transcripts.

### 4.2 Organ-specific *in vivo* Ribo-seq dataset

To provide evidence on the necessity of experiment-specific rRNA depletion, we made use of a comprehensive public dataset that generated *in vivo* Ribo-seq data for multiple tissues in mouse [13]. This dataset includes 9 organs (brain, heart, kidney, liver, lung, pancreas, skeletal muscle, spleen and testis) where experiments are performed in replicates for each organ. The study measures the translation elongation rate differences between different mouse organs with time-course experiments using both harringtonine and cycloheximide. In our analysis we only used samples treated with cycloheximide. One should note that no rRNA depletion protocol was applied within included experiments, however, RNA digestion was performed differently for two groups of samples. For pancreas, spleen and lung, only RNaseT1 nuclease was used for digestion, but for the others, a fixed mix of RNaseT1 and RNaseS7 nucleases was used. This difference enables us to analyze not only the tissue specificity of rRNA fragments but also their technical dependency on the used experimental protocol. Raw sequencing data (fastq files) of this dataset were first downloaded through NCBI with GSE112223 and SRP136268 accession IDs, then, raw reads were trimmed using the cutadapt tool [15] before running the *Ribo-ODDR* pipeline (*design-mode*) with generated trimmed read files.

### 4.3 The *Ribo-ODDR* pipeline

The primary aim of the *Ribo-ODDR* pipeline is to aid the biotinylated oligo design process for the depletion of rRNA fragments in Ribo-seq experiments. This includes designing novel oligos based on pilot experimental data and cross-species optimization of pre-compiled oligo sets. The *Ribo-ODDR* software comes as an executable Python3 script and has a flexible design for different user needs. It does not contain any compiled information on rRNA sequences, enabling *Ribo-ODDR* to be applied on an organism of choice as long as rRNA (or other depletion-intended RNA) sequences are provided by the user.

In the subsections below, we first describe how *Ribo-ODDR* performs the cross-species optimization of pre-compiled oligo sets in its *cross-species optimization mode*. Then, we continue with the details of the used methodology, when designing novel oligos based on the pilot experimental data in the *novel oligo design mode*. Designed oligos, regardless of the *Ribo-ODDR* mode used, are reported to the user in *FASTA, CSV, BED* and *GFF3* file formats. These files contain various relevant information on oligo designs, including depleting potential (percentage of rRNA fragments that can be depleted with that oligo) in pilot samples, positions of the targeted rRNA regions, GC contents, hybridization energies and oligo self-folding statistics. One should also note that, in the *novel oligo design mode, Ribo-ODDR* does not provide a final optimal set of oligos to deplete rRNA fragments. Instead, it reports the depleting potential of all high potential oligos to the user together with other information on these designs. This is in parallel with our flexible software approach. However, *Ribo-ODDR* provides a ‘*Ribo-ODDR oligo-selector*’ user interface to aid the oligo selection process. This interface is presented in the last subsection below. The full workflow of the *Ribo-ODDR* (*novel oligo design mode*) is shown in Figure 7.

**Figure 7:**
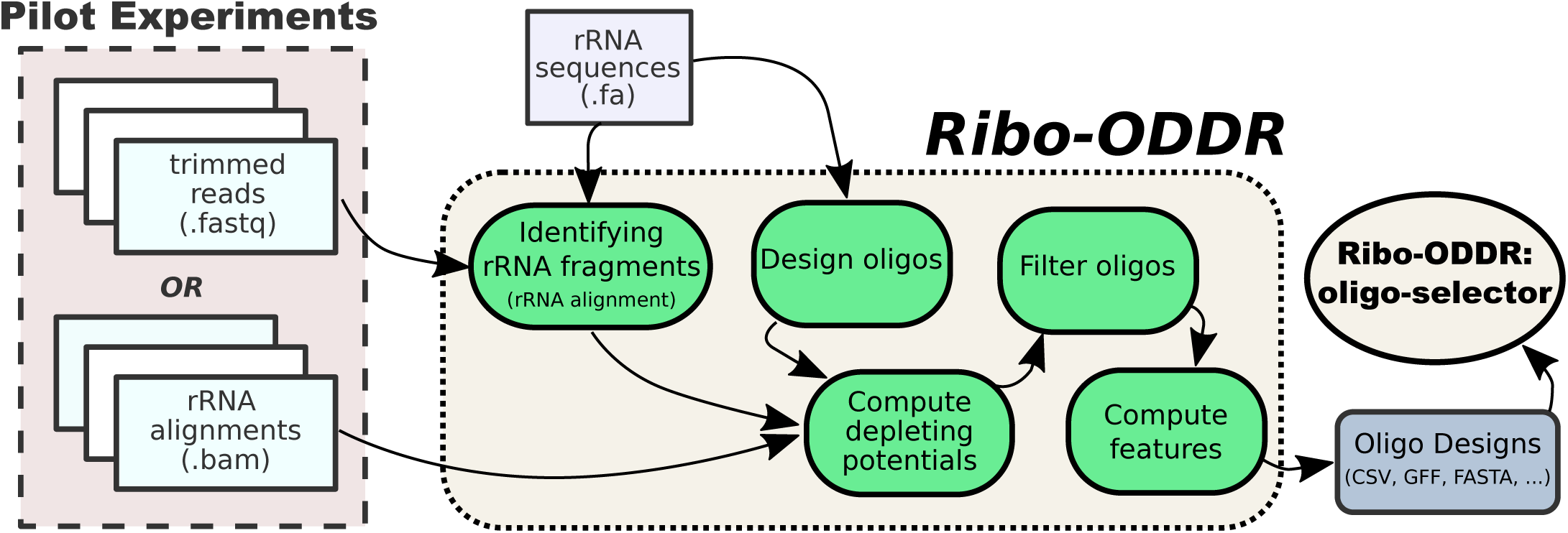
The workflow diagram for the *Ribo-ODDR* pipeline (*novel oligo design mode*).

#### 4.3.1 Cross-species optimization mode

In this mode, *Ribo-ODDR* requires users to provide the sequences of the precompiled oligo set (designed for source organism rRNAs) and target rRNA sequences for the organism to perform Ribo-seq in. Due to the complexity of determining functional homology of rRNA regions between source and target rRNA sequences, we follow a different approach for cross-species optimization. Using *RIsearch2* RNA-RNA interaction prediction tool [18], *Ribo-ODDR* first identifies the most likely target regions of source oligos in the target, accepting the interaction with lowest hybridization energy as most probable. Then, for each source oligo, *Ribo-ODDR* designs the new oligo as perfect complementary to the target interaction region. To reach equivalent coverage as the source oligo, target oligo is later extended on both ends according to oligo dangling ends within the source oligo–target rRNA interaction structure (extending by one for each unpaired nucleotide on 5′ and 3′ ends of the oligo). Note that in the presence of high sequence homology between source and target rRNA sequences, *Ribo-ODDR* can report the same source oligos as optimized oligos.

#### 4.3.2 Novel oligo design mode

The aim of this mode is to compute the depleting potential of all novel oligos on the given pilot Ribo-seq data. In a simple use-case, it requires the user to provide the rRNA sequences, the length range of desired oligos, and the pilot data in which rRNA fragments are abundant. First step of the mode is the identification of rRNA fragments through aligning Ribo-seq reads to user-given rRNA sequences. Next, information on these fragments are used to calculate the depleting potential of all potential oligos that satisfies the user-given length range. Later, *Ribo-ODDR* outputs this information together with various statistics on the designed oligos and final selection of oligos is done by the user using ‘*Ribo-ODDR oligo-selector*’, a user-friendly straightforward R-Shiny user interface.

##### Identifying rRNA fragments

Several variations of the Ribo-seq protocol exist, and for most the generated sequencing data requires trimming of adapter sequences and/or cleaning of used size selection markers before aligning to genome or transcriptome. *Ribo-ODDR* does not perform these pre-processings itself, therefore, requires user to preprocess the sequencing data prior to *Ribo-ODDR*. Under default settings, trimmed & cleaned reads, provided as input pilot data by the user, are first aligned to rRNA sequences using the TopHat aligner [16]. This is done with the following parameter settings, *-n 2 --no-novel-juncs - -no-novel-indels --no-coverage-search --segment-length 25*. However, users can also perform this alignment using other read-aligners and provide the generated bam files as input to *Ribo-ODDR*.

##### Oligo-set generation and depleting potential computation

Next, based on the user-given oligo length range constraint, depletion oligos are generated in a position specific manner. Oligo designs correspond to fixed length regions within user-given rRNA sequences, an oligo sequence being the perfect complementary sequence to its region. Note that the final oligo-set spans all possible regions across all given rRNA sequences. Therefore, oligo designs overlap with each other but the depleting potential of each oligo is computed separately. Following a heuristic approach, *Ribo-ODDR* computes the depleting potential of an oligo (separately for each pilot sample) based on the number of depletable rRNA fragments, i. e. reads that are aligned to the corresponding oligo region within an rRNA. To allow sub-optimal binding between rRNA fragments and the oligo, a fragment (read) is considered depletable only if it satisfies the following constraints. The rRNA fragment has to cover a minimum of 10 nucleotides or two thirds of the oligo length, whichever is higher, within the oligo region under consideration. Additionally, the rRNA fragment can have a maximum of 10 nucleotides or one third of the oligo length, whichever is lower, outside the oligo region to be considered as depletable by that oligo. Based on these constraints, for each pilot sample, we simply count the number of depletable fragments for every oligo and report its percentage within all rRNA fragments as the depleting potential for each pilot sample.

##### Filtering oligos based on depleting potential

For fast computation of oligo features, *Ribo-ODDR* filter outs some of the low potential oligos based on customizable thresholds. Under default settings, it discards the oligos that have a depleting potential less than 0.05 (5% of all rRNA fragments) in more than 75% of the provided pilot samples. However, these thresholds can be altered by the user.

##### Computation of other oligo features

In addition to sample-specific depleting potential of oligos in pilot samples, *Ribo-ODDR* reports a few other informative statistics on designed oligos, for which some are straightforward like *GC content* and *target rRNA position*. For each oligo, an *over-all depletion score* is also computed by *Ribo-ODDR*, that is the ratio of samples oligo has a depleting potential more than a user-given threshold, 0.05 (5%) in default settings. Additionally, for each oligo, *Ribo-ODDR* reports a *minimum hybridization energy* that is the free energy of the full perfect compli-mentary binding to an rRNA fragment at 37°C computed by *RIsearch2* [18]. Using the *RNAfold* program from the ViennaRNA Package [19], self-folding of the oligo is also predicted. This is reported in three different features, predicted *structure*, the *MFE* as the free energy of the predicted structure, and the *base pairing percentage* within the given structure.

##### Off-target prediction for designed oligos

If protein-coding transcript sequences of the organism are provided by the user, *Ribo-ODDR* computes the off-targeting potential of oligos as well. Denoting the minimum binding free energy across all oligos as *E*_*min*_ and the minimum oligo length as *l*_*min*_, oligo off-targets are predicted using *RIsearch2* [18] with the following parameter settings, 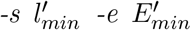, where 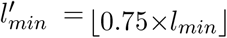 and 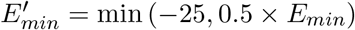. These settings allow us to detect the potential off-target regions on given transcripts, that have a considerably low binding energy with designed oligos. Number of predicted off-targets are reported to the user as an additional oligo feature, however, the additional information on individual off-target predictions are outputted separately.

#### 4.3.3 Selecting final oligos with *Ribo-ODDR oligo-selector*

To aid the final selection of oligos from all oligos outputted by the *Ribo-ODDR novel oligo design mode*, we present the *Ribo-ODDR oligo-selector* auser interface, that uses the R-shiny environment. In this interface, users can explore the features of the designed oligos within the available-oligo-list, filter them according to different filters on reported features and add the desired ones to the selection list, which results in removing the overlapping oligos from the available oligo list. A snapshot from this interface is shown in Supplementary Figure 16.

### 4.4 Experimental details on Ribo-seq experiments

*C57BL/6* female and male mice between 8 and 12 weeks of age were used for experiments. For Figure 5, both *Lgr5Cre*^*ERT2*^ [20] *and VillinCre*^*ERT2*^ [21] *mice were crossed to the RiboTag mouse [22] to generate Lgr5Cre*^*ERT2*^*-RPL22*.*HA* and *VillinCre*^*ERT2*^*-RPL22*.*HA* respectively. Due to differences in recombination efficiency and total number of cells, the tamoxifen-mediated induction of Cre-recombinase varied slightly between the two lines: for the *Lgr5Cre*^*ERT2*^*-RPL22*.*HA* mice, recombination was induced by a single intraperitoneal injection of 120 mg/kg tamoxifen and samples were taken for downstream analysis after 24h and 48h; for the *VillinCre*^*ERT2*^*-RPL22*.*HA* mice, recombination was induced after two consecutive intraperitoneal injections of 80 mg/kg tamoxifen and samples were taken after 120h. Due to availability of strains, for Figure 6, *Lgr5Cre*^*ERT2*^*-RPL22*.*HA* and *Lgr5Cre*^*ERT2*^*-Rptor*^*fl/fl*^ mice were crossed to generate *Lgr5Cre*^*ERT2*^*-Rptor*^*fl/fl*^*-RPL22*.*HA* animals. Recombination was induced by a single intraperitoneal injection of 120 mg/kg tamoxifen and samples were taken for downstream analysis after 24h. Mice were bred in-house at the Netherlands Cancer Institute and all experimental protocols were approved by the NKI Animal Welfare Body.

#### 4.4.1 Sample preparation from *in vivo* small intestines

Mice were euthanized by CO_2_ and small intestines were immediately dissected, flushed with cold PBS supplemented with 100 µg/mL of cycloheximide and snap frozen using liquid nitrogen. Frozen tissues were ground by pestle and mortar while submerged in liquid nitrogen. The resulting powder was rapidly dissolved in cold lysis buffer (20 mM Tris HCl pH 7.4, 10 mM MgCl_2_, 150 mM KCl, 1% NP-40, 100 µg/mL cycloheximide and 1x EDTA-free proteinase inhibitor cocktail (Roche, 04693132001)) and incubated on ice for 30min. Samples were then homogenized using a Tissue Lyser (3 rounds of 45 sec at 50 oscillations per second) and centrifuged at max speed for 20min at 4 °C.

#### 4.4.2 Sample preparation from *in vitro* crypt cultures

Crypt cultures were generated from the *VillinCre*^*ERT2*^*-RPL22*.*HA* mice as described previously [23]. Around 120 plugs of 30 µL BME (Amsbio #3533-010-02) were used for each sample. Ribosomes were stalled by incubating cells with 100 µg/mL cycloheximide for 3-5min at 37 °C, after which all steps were carried on ice. Cells were collected and washed twice in cold PBS supplemented with 100 µg/mL cycloheximide, and homogenized with a 25G needle in cold lysis buffer. After incubating the lysates on ice for 20min, samples were centrifuged at max speed for 20min at 4 °C.

#### 4.4.3 Ribosome profiling

##### Pull down of HA-tag ribosomes

All supernatants (from both *in vivo* small intestines and *in vitro* crypt cultures) were pre-cleared for 20min at 4 °C, using Pierce™ Control Agarose Matrix (ThermoFisher #26150), after which they were incubated with pre-washed Anti-HA.11 Epitope Tag Affinity Matrix (BioLegend #900801) overnight at 4 °C. Ribosomes were eluted in lysis buffer containing 200 µg/mL HA peptide (ThermoFisher #26184) and supplemented with 100 µg/mL cycloheximide for 15min at 30 °C. Exposed RNA was digested with RNase I (ThermoFisher #AM2294) for 40min at 25 °C and this process was stopped by adding SUPERASE (ThermoFisher #AM2694). RPFs were purified using the miRNeasy minikit (Qiagen #217004) following the manufacturer’s protocol and used for the library preparation.

##### Library preparation

The library preparation was conducted as previously described [14] with some modifications. Briefly, RPFs were run in a 10% TBE-Urea polyacrylamide gel and size selected between 19 nt and 32 nt as marked by RNA oligonucleotides (see Supplementary Table 5). Gel slices were crushed, eluted and ethanol precipitated. Samples were then dephosphorylated in the 3′ region using T4 polynucleotide kinase (PNK) (NEB #M0201) and 1.5xMES buffer (150 mM MES-NaOH, 15 mM MgCl_2_, 15 mM β-mercaptoethanol and 450 mM NaCl, pH 5.5) and incubated at 37 °C for 4h. RNAs were purified using Tri-zol and the 3′ adapter (see Supplementary Table 5) was added using T4 RNA ligase I (NEB #M0204) at 24 °C overnight. The ligated products were size selected and 5′ phosphorylated with T4 PNK for 30min at 37 °C. After purifying the RNAs, the 5′ adaptor (see Supplementary Table 5) was added with T4 RNA ligase I for 2,5h at 37 °C and the final products with both adaptors were size selected one last time on a 10% TBE-Urea polyacrylamide gel. This was followed by rRNA depletion, which was performed according to manufacturer’s instructions RiboZero (Illumina, catalog no. 20020598) and RiboCop (Lexogen, catalog no. 037) here) when using commercial kits. For rRNA depletion with custom oligos, samples were incubated with 2 µL of the different biotinylated oligos (10 µM each oligo, Supplementary Table 1-4) in 20 µL with 2xSSC (ThermoFisher #15557044). Samples were then denatured at 100 °C for 1min, followed by an incubation at 37 °C for 15min. In the meantime, 40 µL of MyOne Streptavidin C1 DynaBeads (ThermoFisher #65001) were washed and re-suspended in 20 µL of 2x wash/bind buffer (2 M NaCl, 1 mM EDTA, 5 mM Tris and 0.2% Triton X-100) and mixed with the sample at 1000rpm for 30min and at 37 °C. Supernatants were collected and RNAs were precipitated with isopropanol and resuspended in 8 µL of RNase-free water. Reverse transcription was performed with SuperScript III (ThermoFisher #18080051) following the manufacturer’s instructions and using the RTP primer (see Supplementary Table 5). cDNA was then purified using G-50 columns (Merck GE27-5330-01) and used as a template for the PCR reaction with Phusion High-Fidelity DNA Polymerase (ThermoFisher #F530L) for 18 cycles, with primers listed in Supplementary Table 5. PCR products were purified using the QIAquick PCR purification kit (Qiagen #28104) followed by a E-Gel SizeSelect II 2%, (ThermoFisher #G661012). The quality and molarity of the samples were evaluated with the Agilent 2100 Bioanalyzer and the libraries were sequenced on the Illumina HiSeq2500 by the Genomics Core Facility at the Netherlands Cancer Institute.

##### Data processing

Raw reads are trimmed and cleaned from the size selection markers using the cutadapt tool [15]. Then, *Ribo-ODDR* (*design-mode*) is run with generated trimmed read files to align reads to mouse rRNA sequences (28S: NR 003279.1, 18S: NR 003278.3, 5.8S: NR 003280.2, 5S: NR 030686.1) and to design depletion oligos. Preprocessed reads are cleaned from rRNA fragments using the *SortMeRNA* tool [17] and remaining reads are mapped to gencode release M21 protein-coding transcript sequences and GRCm38.p6 (mm10) mouse genome using the TopHat aligner [16].

## 5 CONCLUSION

In this study, we show that the use of commercial rRNA depletion kits may result in suboptimal depletion in Ribo-seq experiments, and that different tissues and experimental conditions result in heterogeneity of produced rRNA fragments. Both of these findings demonstrate the necessity of experiment-specific custom oligo design for efficient rRNA depletion. To aid the computational part of the oligo design process, we have developed *Ribo-ODDR*, a Ribo-seq focused oligo design pipeline for experiment-specific rRNA depletion. Oligos designed using this platform resulted in a substantial increase in rRNA depletion *in vivo* Ribo-seq experiments in mouse intestine, with much higher depletion performance when compared to commercial kits. Ultimately, this allows higher sequencing depth on the translatome and more powerful down-stream analyses of the data. The tool is easy to use, and will allow the optimization of this crucial step in the Ribo-seq protocol, particularly for samples that have proven difficult to assay. *Ribo-ODDR* is an open source software and freely accessible at https://github.com/fallerlab/Ribo-ODDR.

## 6 ACKNOWLEDGEMENTS

We would like to thank past and present members of the Faller Lab and Abhijeet Pataskar for critical reading and fruitful discussions.

## Supplementary Figures and Tables

- **Supplementary Figure 1:** Analysis of an additional mouse Ribo-seq dataset generated using Ribo-Zero.
- **Supplementary Figure 2:** Suboptimal performance of RiboCop rRNA depletion kit in Ribo-seq experiment using *in vitro* mouse intestinal organoids.
- **Supplementary Figure 3:** Sample-specificity of rRNA fragments and cross-replicate correlation analysis of oligo depleting potentials for brain.
- **Supplementary Figure 4:** Sample-specificity of rRNA fragments and cross-replicate correlation analysis of oligo depleting potentials for heart.
- **Supplementary Figure 5:** Sample-specificity of rRNA fragments and cross-replicate correlation analysis of oligo depleting potentials for kidney.
- **Supplementary Figure 6:** Sample-specificity of rRNA fragments and cross-replicate correlation analysis of oligo depleting potentials for liver.
- **Supplementary Figure 7:** Sample-specificity of rRNA fragments and cross-replicate correlation analysis of oligo depleting potentials for skeletal muscle.
- **Supplementary Figure 8:** Sample-specificity of rRNA fragments and cross-replicate correlation analysis of oligo depleting potentials for testis.
- **Supplementary Figure 9:** Sample-specificity of rRNA fragments and cross-replicate correlation analysis of oligo depleting potentials for lung.
- **Supplementary Figure 10:** Sample-specificity of rRNA fragments and cross-replicate correlation analysis of oligo depleting potentials for pancreas.
- **Supplementary Figure 11:** Sample-specificity of rRNA fragments and cross-replicate correlation analysis of oligo depleting potentials for spleen.
- **Supplementary Figure 12:** Tissue and RNase specificity of rRNA fragments in mouse, based on positional abundance profile of 18S, 5-8S and 5S rRNA fragments.
- **Supplementary Figure 13:** rRNA abundance profiles of *in vitro* Ribo-seq experiments with human (SET-0) and SET-1 oligos.
- **Supplementary Figure 14:** Alternative visualisation for rRNA abundance profiles of *in vivo* Ribo-seq experiments with SET-1 and SET-2 oligos.
- **Supplementary Figure 15:** Read count analysis for potential off-targets of new SET2 oligos.
- **Supplementary Figure 16:** Screenshot from the oligo selection user interface, *Ribo-ODDR oligo-selector*.
- **Supplementary Table 1:** Sequences for human oligos (SET-0).
- **Supplementary Table 2:** Sequences for SET-1 oli-gos.
- **Supplementary Table 3:** Sequences for SET-2 oli-gos.
- **Supplementary Table 4:** Sequences for SET-3 oli-gos.
- **Supplementary Table 5:** adapter and primer sequences.

