## Supplementary Material for "*Ribo-ODDR*: Oligo Design pipeline for experiment-specific Depletion of Ribosomal RNAs in Ribo-seq"

### *Ribo-ODDR*: *O*ligo *D*esign pipeline for experiment-specific *D*epletion of *R*ibosomal RNAs in *Ribo*-seq
(Supplementary Document)

Ferhat Alkan [†,*], Joana Silva [†], Eric Pintó Barberà, William J Faller [*]

Division of Oncogenomics, The Netherlands Cancer Institute, 1066 CX, Amsterdam, The Netherlands

[†] Joint first authors

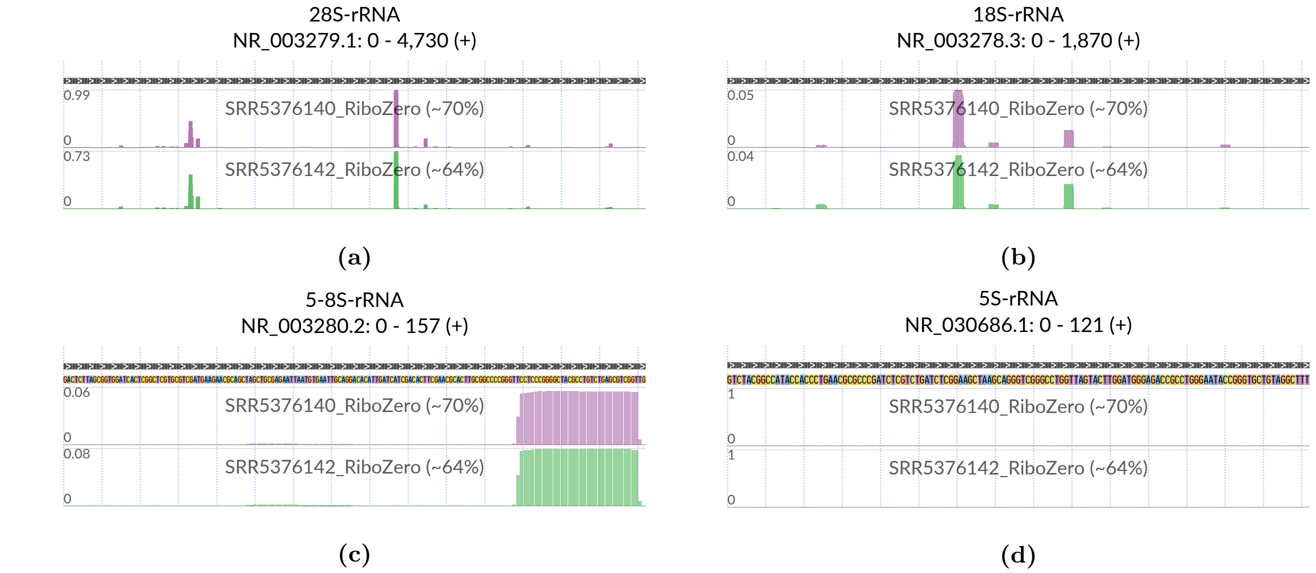

**Supplementary Figure 1:** Suboptimal performance of Ribo-Zero rRNA depletion kit in Ribo-seq. Visualization is based on public data[1] where Ribo-Zero kit was used for rRNA depletion. Each track shows the positional abundance profile of 28S (a), 18S (b), 5-8S (c) and 5S (d) rRNA fragments coming from individual samples. For every position in the x-axis, y-axis represents the normalized read ratio, number of rRNA reads mapped to that position divided by the total number of reads mapped to all protein coding transcripts. Sample-specific total rRNA percentages are given in track labels together with public SRA accession IDs for analyzed experiments.

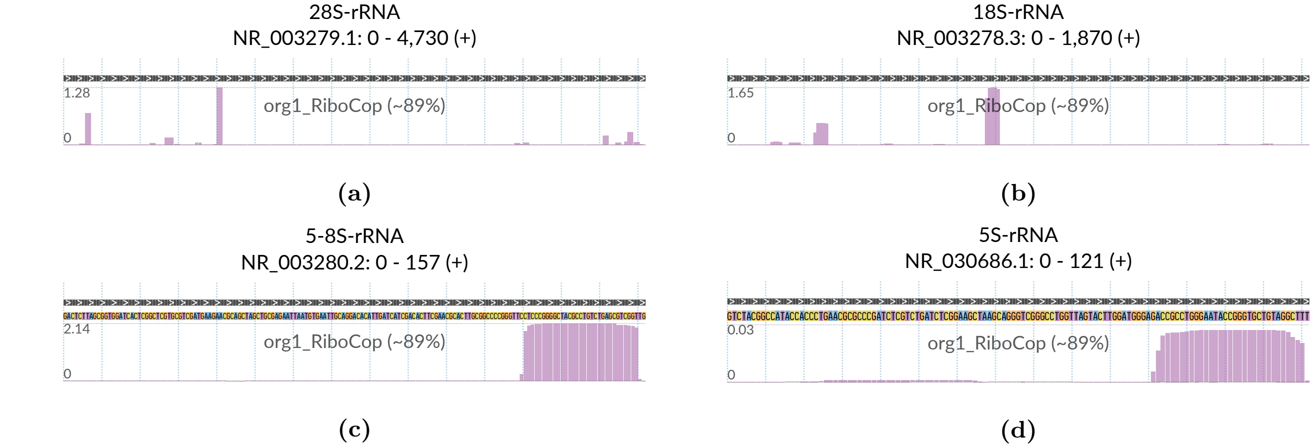

**Supplementary Figure 2:** Suboptimal performance of RiboCop rRNA depletion kit in Ribo-seq. Visualization is based on an *in vitro* Ribo-seq experiment using RiboCop, performed in mouse intestinal organoids. The track shows the positional abundance profile of 28S (a), 18S (b), 5-8S (c) and 5S (d) rRNA fragments. For every position in the x-axis, y-axis represents the normalized read ratio, number of rRNA reads mapped to that position divided by the total number of reads mapped to all protein coding transcripts. Total rRNA percentage is given in the track label.

[1] Simsek D, Tiu GC, Flynn RA, Byeon GW, Leppek K, Xu AF, Chang HY, Barna M. The Mammalian Ribo-interactome Reveals Ribosome Functional Diversity and Heterogeneity. Cell. 2017 Jun; 169(6):1051–1065.

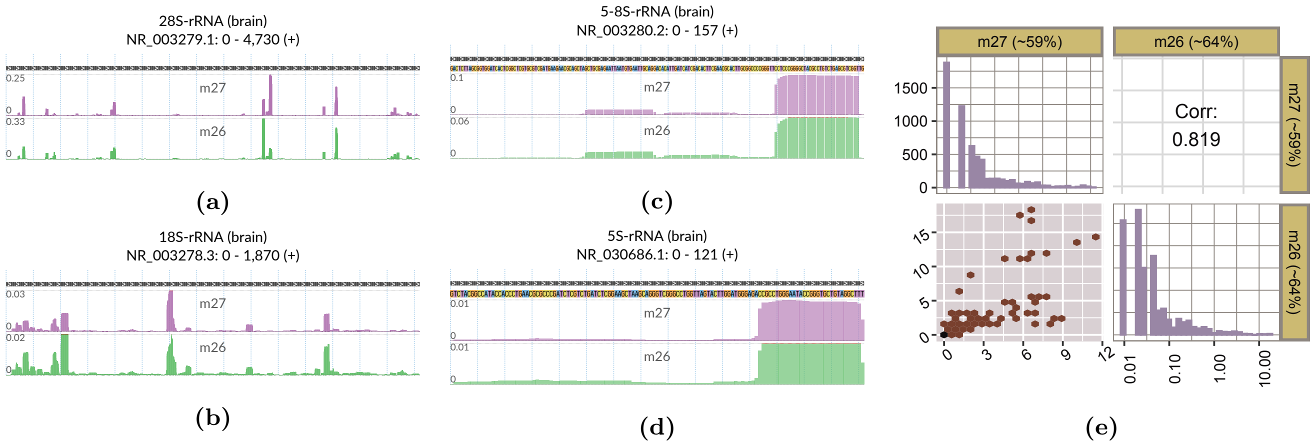

**Supplementary Figure 3:** Sample-specific rRNA fragment profiles (a-d) of **brain** replicates and the cross-replicate correlation analysis (e) of oligo depleting potentials computed by *Ribo-ODDR*. These figures are generated in a similar way to Figure 2&3 of the original paper, with the exception of being sample-specific over all **brain** replicates instead of being organ-specific at each row and column.

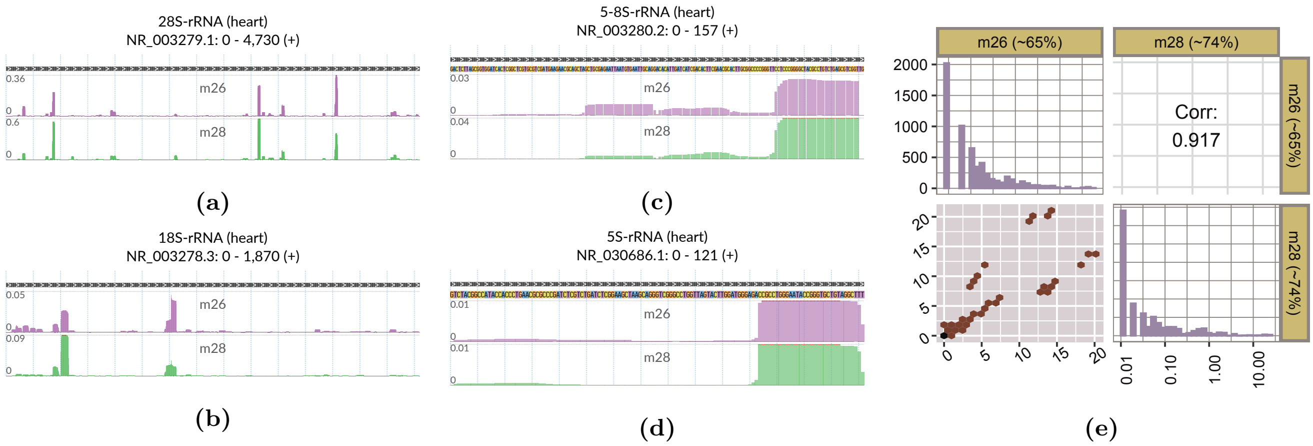

**Supplementary Figure 4:** Sample-specific rRNA fragment profiles (a-d) of **heart** replicates and the cross-replicate correlation analysis (e) of oligo depleting potentials computed by *Ribo-ODDR*. These figures are generated in a similar way to Figure 2&3 of the original paper, with the exception of being sample-specific over all **heart** replicates instead of being organ-specific at each row and column.

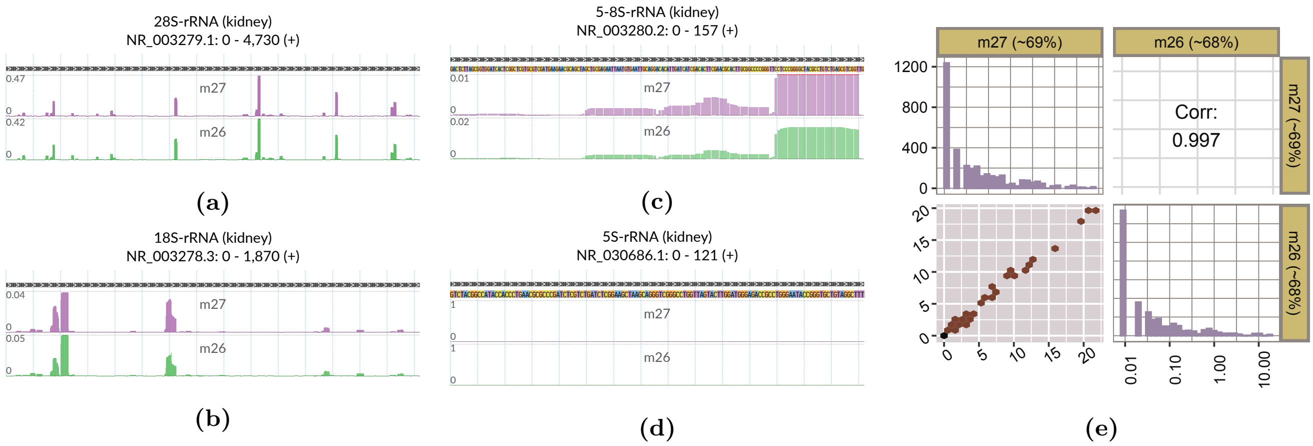

**Supplementary Figure 5:** Sample-specific rRNA fragment profiles (a-d) of **kidney** replicates and the cross-replicate correlation analysis (e) of oligo depleting potentials computed by *Ribo-ODDR*. These figures are generated in a similar way to Figure 2&3 of the original paper, with the exception of being sample-specific over all **kidney** replicates instead of being organ-specific at each row and column.

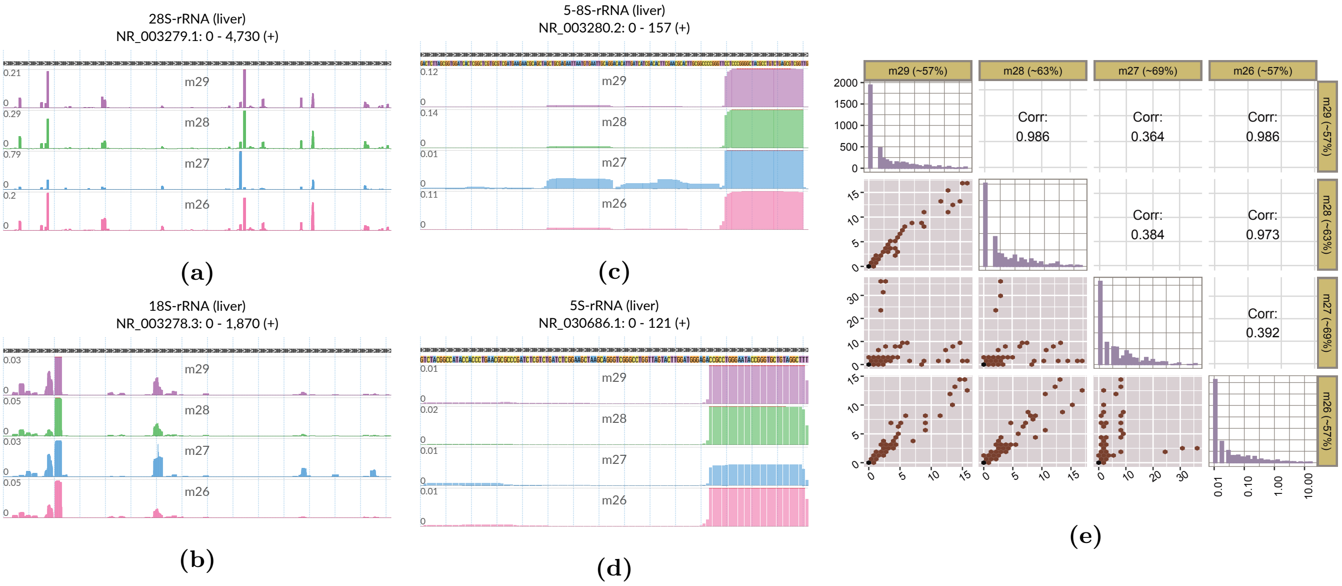

**Supplementary Figure 6:** Sample-specific rRNA fragment profiles (a-d) of **liver** replicates and the cross-replicate correlation analysis (e) of oligo depleting potentials computed by *Ribo-ODDR*. These figures are generated in a similar way to Figure 2&3 of the original paper, with the exception of being sample-specific over all **liver** replicates instead of being organ-specific at each row and column.

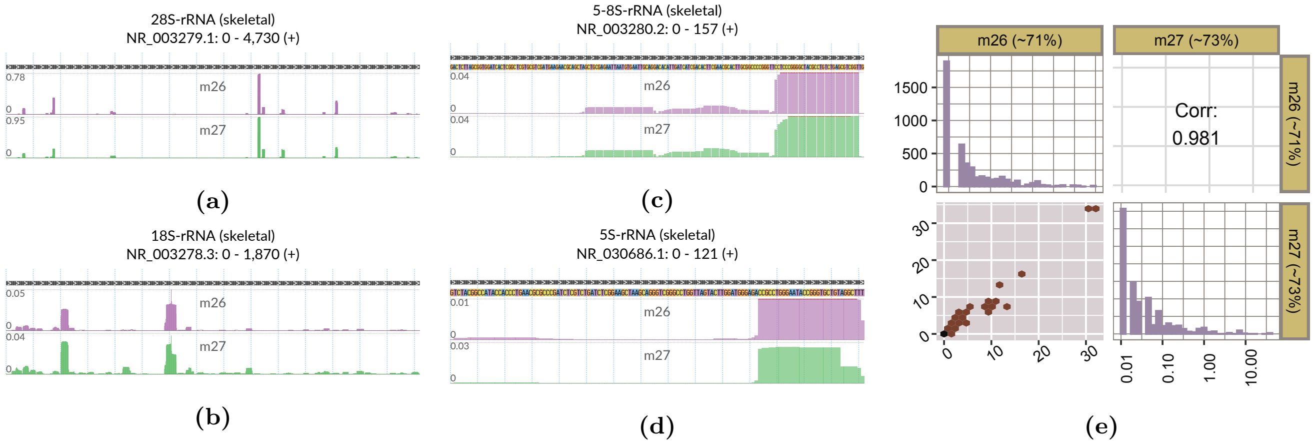

**Supplementary Figure 7:** Sample-specific rRNA fragment profiles (a-d) of **skeletal** replicates and the cross-replicate correlation analysis (e) of oligo depleting potentials computed by *Ribo-ODDR*. These figures are generated in a similar way to Figure 2&3 of the original paper, with the exception of being sample-specific over all **skeletal** replicates instead of being organ-specific at each row and column.

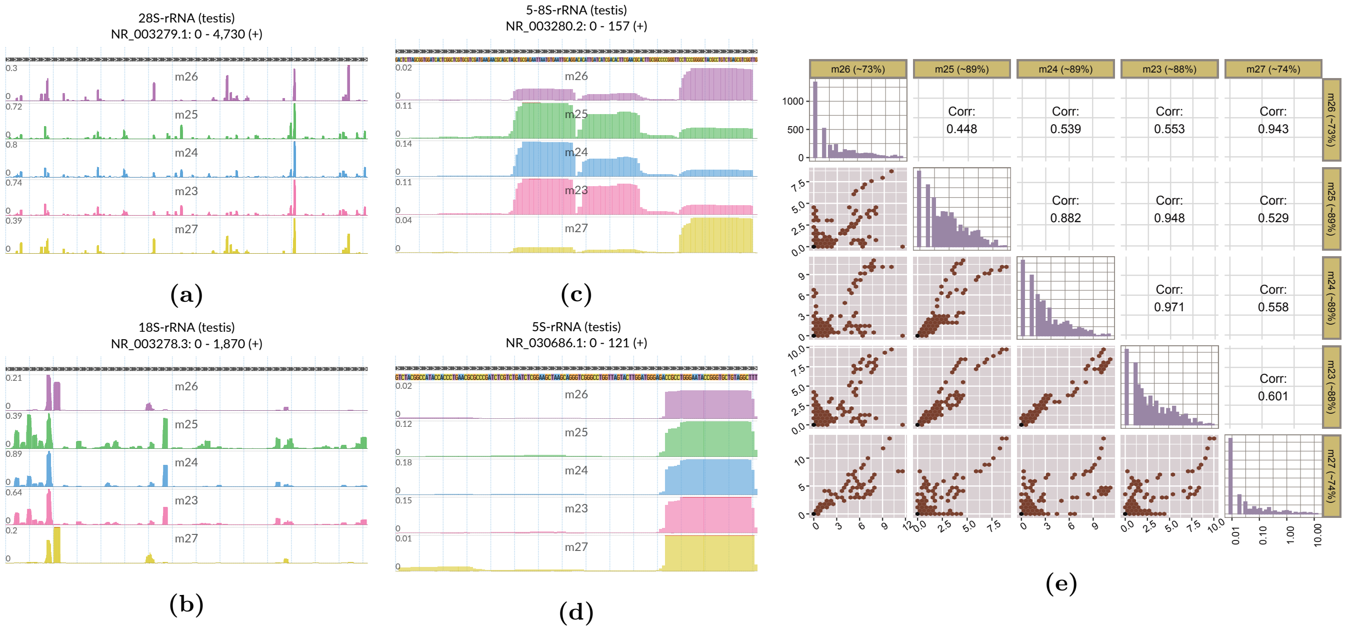

**Supplementary Figure 8:** Sample-specific rRNA fragment profiles (a-d) of **testis** replicates and the cross-replicate correlation analysis (e) of oligo depleting potentials computed by *Ribo-ODDR*. These figures are generated in a similar way to Figure 2&3 of the original paper, with the exception of being sample-specific over all **testis** replicates instead of being organ-specific at each row and column.

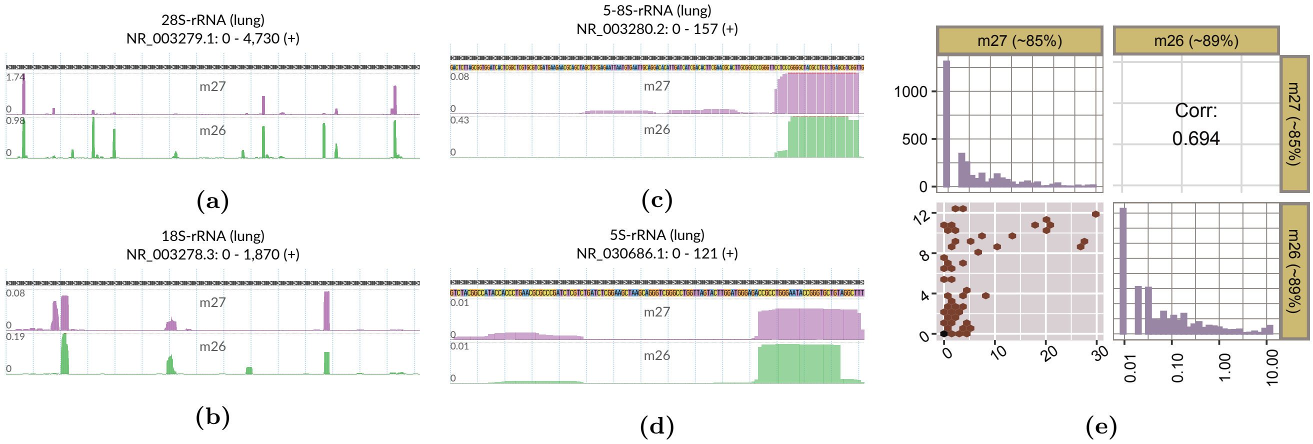

**Supplementary Figure 9:** Sample-specific rRNA fragment profiles (a-d) of **lung** replicates and the cross-replicate correlation analysis (e) of oligo depleting potentials computed by *Ribo-ODDR*. These figures are generated in a similar way to Figure 2&3 of the original paper, with the exception of being sample-specific over all **lung** replicates instead of being organ-specific at each row and column.

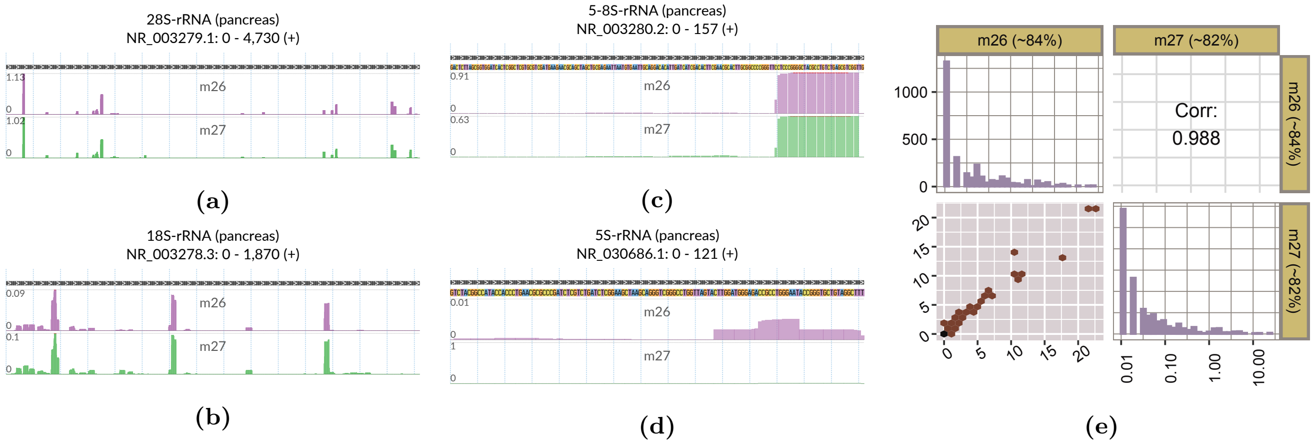

**Supplementary Figure 10:** Sample-specific rRNA fragment profiles (a-d) of **pancreas** replicates and the cross-replicate correlation analysis (e) of oligo depleting potentials computed by *Ribo-ODDR*. These figures are generated in a similar way to Figure 2&3 of the original paper, with the exception of being sample-specific over all **pancreas** replicates instead of being organ-specific at each row and column.

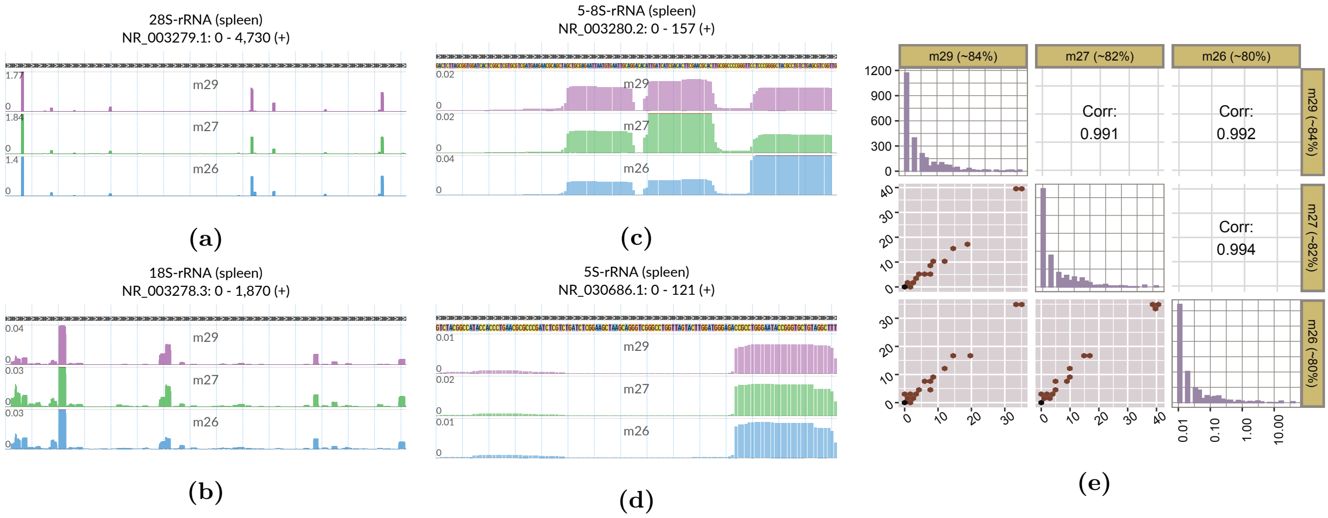

**Supplementary Figure 11:** Sample-specific rRNA fragment profiles (a-d) of **spleen** replicates and the cross-replicate correlation analysis (e) of oligo depleting potentials computed by *Ribo-ODDR*. These figures are generated in a similar way to Figure 2&3 of the original paper, with the exception of being sample-specific over all **spleen** replicates instead of being organ-specific at each row and column.

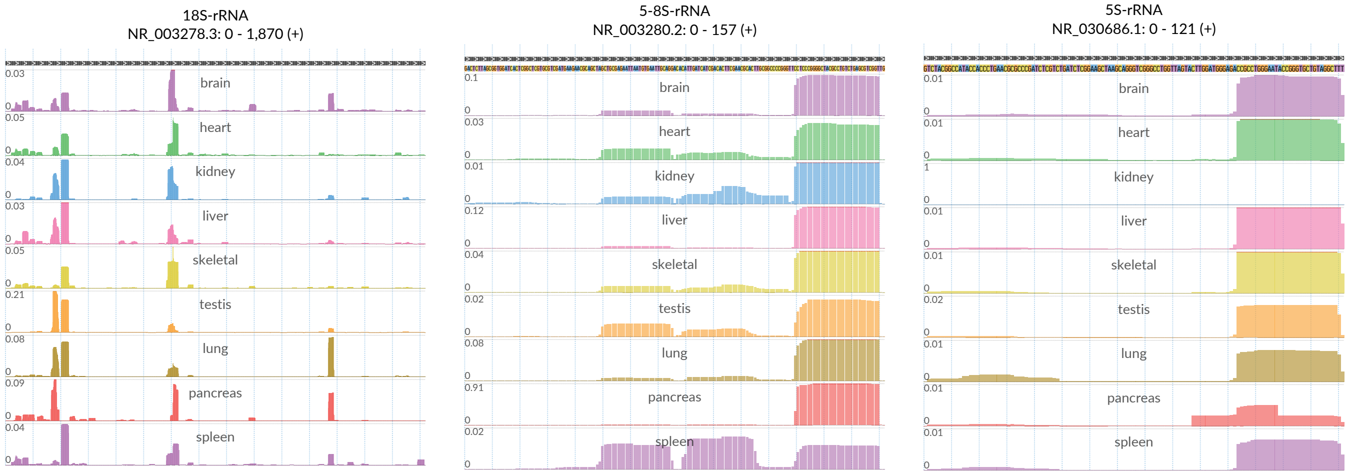

**(a)** **(b)** **(c)**

**Supplementary Figure 12:** Tissue and RNase specificity of rRNA fragments in mouse, based on positional abundance profile of 18S (a), 5-8S (b) and 5S (c) rRNA fragments. For every position in the x-axis, y-axis represents the normalized read ratio, number of rRNA reads mapped to that position divided by the total number of reads mapped to all protein coding transcripts.

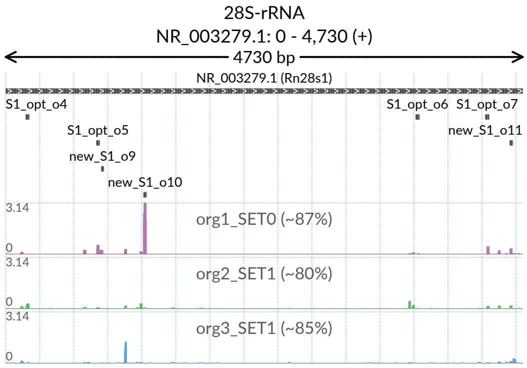

**(a)**

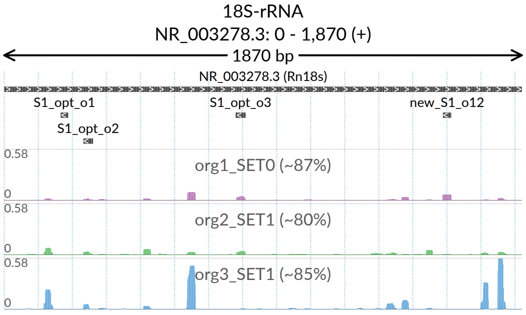

**(b)**

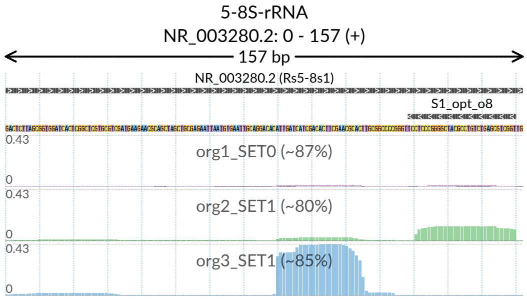

**(c)**

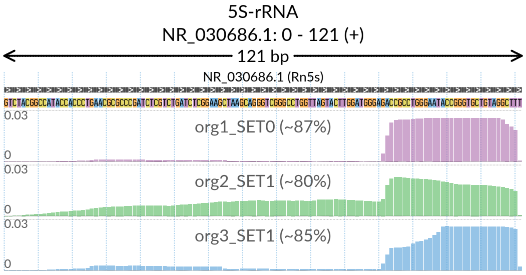

**(d)**

**Supplementary Figure 13:** Positional abundance profiles of 28S (a), 18S (b), 5-8S (c) and 5S (d) rRNA fragments coming from preliminary *in vitro* Ribo-seq experiments performed with two different sets of rRNA depletion oligos (SET-0 (human oligos) & SET-1). Prior to *in vivo* experiments (in mouse intestine) presented in the paper, we first performed an *in vitro* Ribo-seq (in mouse intestinal organoid) using human rRNA depletion oligos (SET-0, Supplementary Table 1). In this experiment, we observed that ∼87% of the reads that mapped to rRNAs and protein-coding transcripts were rRNA fragments and some of these fragments originated from mouse rRNA regions that were actually targeted by human oligos (top row, *org1_SET0*). To improve the overall quality of this experiment, we used the early versions of *Ribo-ODDR cross-species optimization mode* to optimize the human oligos for mouse experiments, and added four new oligos to the pool by manually visualizing rRNA fragments and selecting the hotspots requiring depletion. At the end, the new set of oligos (SET-1) included cross-species optimized version of human oligos together with 4 additional ones (Supplementary Table 2). While this approach did result in increased rRNA depletion, in average only ∼18% of sequencing reads mapped to protein coding regions (bottom 2 rows, *org2_SET1* & *org3_SET1*). In the figure, top track indicates the target regions of used oligos within a rRNA, where additional oligos of the SET-1 are labeled as ‘new’. In all tracks, x-axis corresponds to position within rRNAs. In sample-specific profile tracks, y-axis is fixed to the same interval and shows the normalized read ratio, number of rRNA reads mapped to the position divided by the total number of reads mapped to all protein coding transcripts. The percentages given within sample labels indicates the sample-specific percentage of rRNA fragments, within all reads that can be mapped to rRNA and protein-coding transcripts.

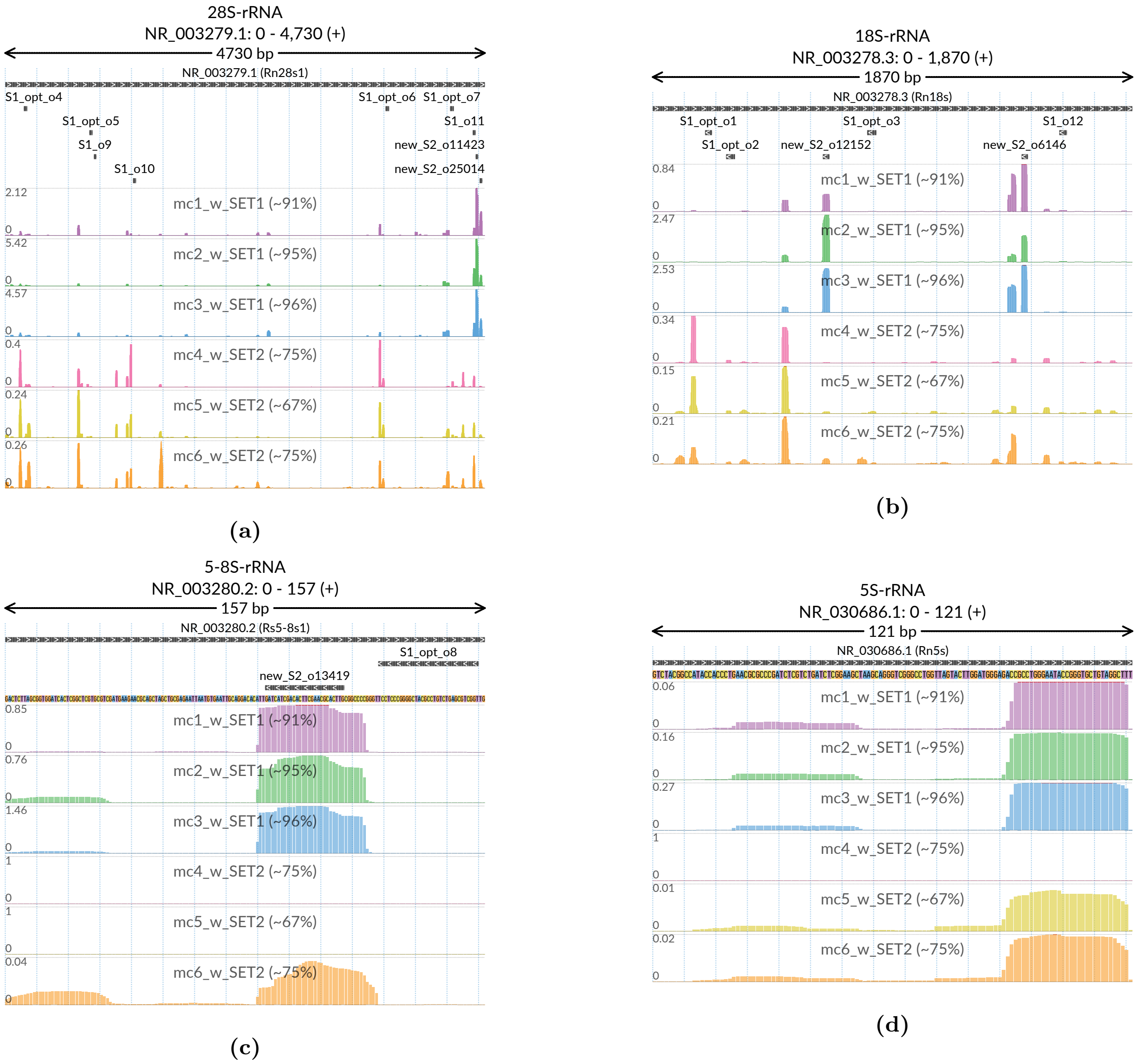

**Supplementary Figure 14:** Positional abundance profiles of 28S (a), 18S (b), 5-8S (c) and 5S (d) rRNA fragments coming from *in vivo* (mouse intestine) Ribo-seq experiments performed with two different sets of rRNA depletion oligos (SET-1 & SET-2). SET-2 includes all oligos of the SET-1 together with 5 additional ones, designed with *Ribo-ODDR* based on pilot data generated with experiments using SET-1. In each figure, top track indicates the target regions of used oligos within that rRNA, where additional oligos of the SET-2 are labeled as 'new'. In all tracks, x-axis corresponds to position within rRNAs. In sample-specific profile tracks, y-axis shows the normalized read ratio, number of rRNA reads mapped to the position divided by the total number of reads mapped to all protein coding transcripts. Y-axis of each row has a different interval between 0 and the max value given in every row. The percentages given within sample labels indicates the sample-specific percentage of rRNA fragments, within all reads that can be mapped to rRNA and protein-coding transcripts.

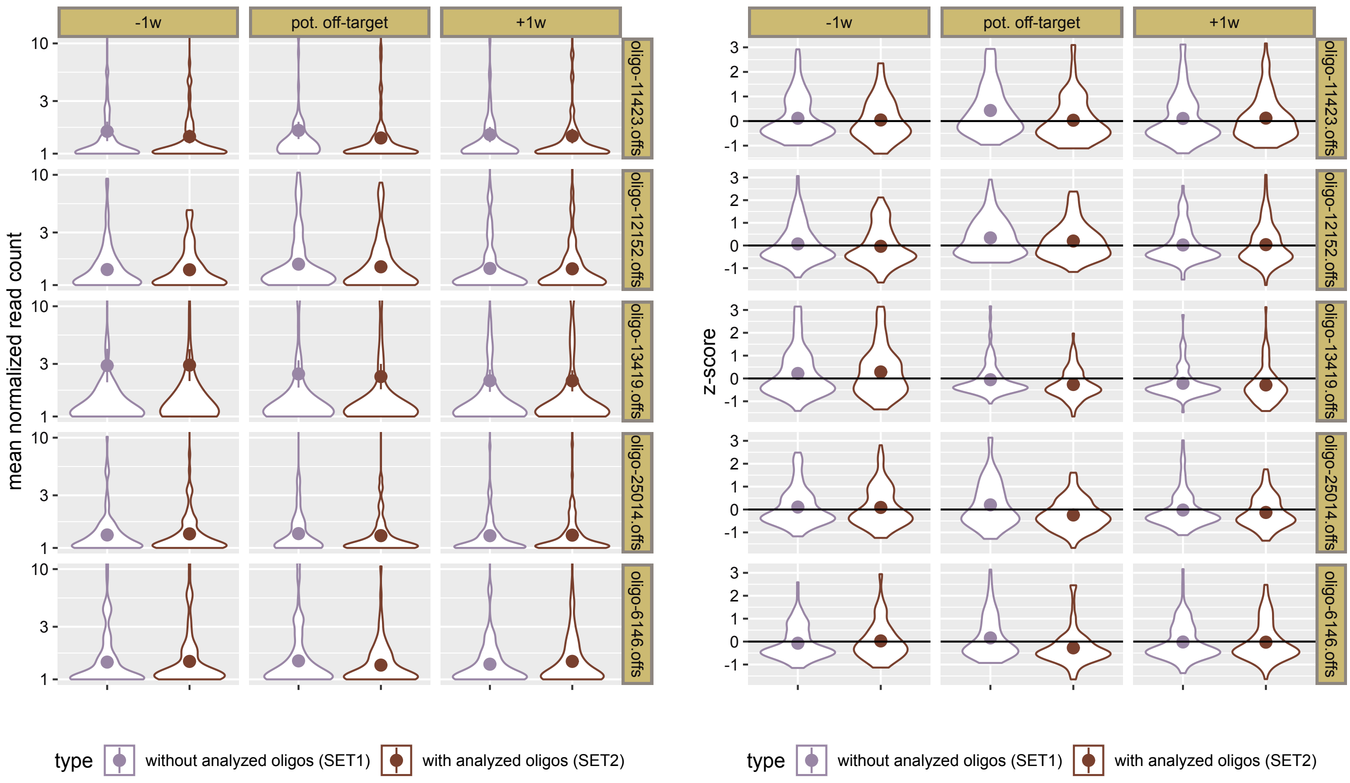

(a) (b)

**Supplementary Figure 15:** Read count analysis for potential off-targets of new SET2 oligos. Potential off-targets of 5 new oligos are first identified with *Ribo-ODDR* & RIsearch2[2] tools, choosing the 5′UTR+CDS of gencode vM21 protein-coding transcripts as unintended depletion targets. To avoid multiple representation of predicted off-target regions due to alternative isoforms, only predictions within primary transcripts are kept. Later, predicted off-target regions, their upstream and downstream regions (5 x off-target region length) are queried for active ribosome protected fragments (RPFs). If no RPF is found in these regions, that off-target is considered not translated and discarded from analysis. All remaining off-target predictions are ranked based on minimum free energy, representing the likelihood of undesired off-target depletion in the suspected region. From this ranking, we select the top 100 off-target predictions for each oligo to perform the read-count analysis. In the analysis, for each of the 500 selected off-target prediction, we first determine the mean normalized read count mapped to each off-target region and their 5 upstream and 5 downstream neigboring regions (same length as the off-target). This is done separately for *in vivo* Ribo-seq experiments performed with SET-1 and SET-2 oligos. This yields to 11 values per oligo off-target prediction and experiment group, each representing the mean normalized read count of a region (off-target or neigbor) within the experiments using one of the two oligo sets. We store these values not only as it is but also by transforming them into z-scores based on the distribution of these 11 values. In the figure above, on the left (a), we first present how original mean normalized read counts are distributed for SET-1 and SET-2 experiments (different colors), within off-target regions and their immediate neigbors (columns), seperately for each oligo analyzed (rows). The candidate region for unintended depletion is highlighted with red lines and neighboring regions serve as control. On the right (b), instead of original scores, we present, in the same way, the distribution of z-scores, calculated seperately for each oligo off-target prediction and experiment group. In both figures, one can see that reads within the predicted off-target regions (center column) do not deplete due to unintended oligo targeting, having similar distributions between SET-1 and SET-2 experiments, just like as expected for their neigboring regions (left and right columns).

[2]Alkan F, Wenzel A, Palasca O, Kerpedjiev P, Rudebeck AF, Stadler PF, Hofacker IL, Gorodkin J. RIsearch2: suffix array-based large-scale prediction of RNA-RNA interactions and siRNA off-targets. Nucleic Acids Res. 2017 05; 45(8):e60.

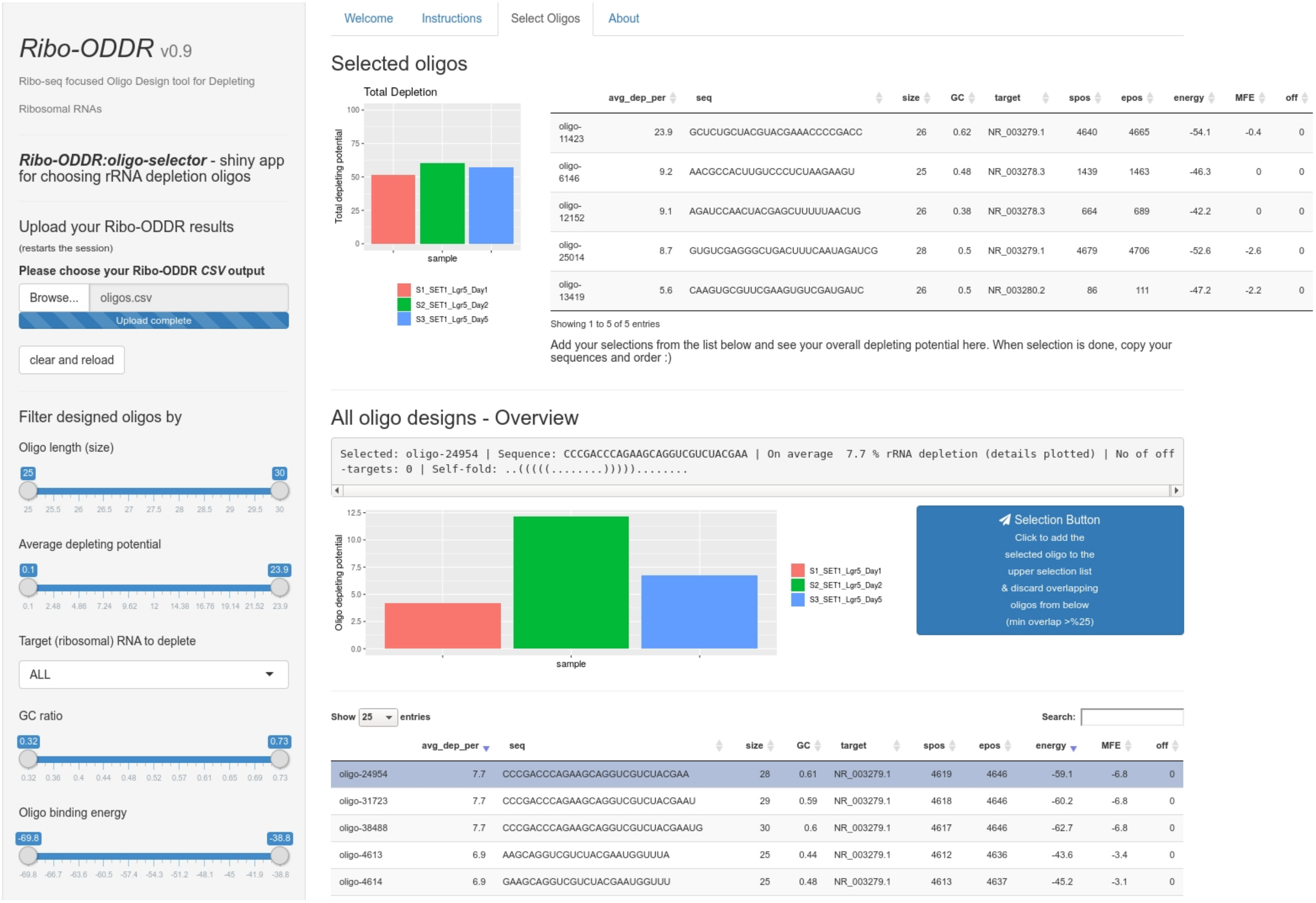

**Supplementary Figure 16:** Screenshot from the oligo selection user interface, *Ribo-ODDR oligo-selector*.

| Oligo ID | sequence (5′ –> 3′) | target rRNA |
|---|---|---|
| human-o1 | UGAUCUGAUAAAUGCCGCAUCCCCCC | 18S |
| human-o2 | CGUGCGAUCGGCCCGAGGUUAUCUAGAGUCACCAA | 18S |
| human-o3 | AUUCCAUUAUUCCUAGCUGCGGUAUCCAGGCGGCUC | 18S |
| human-o4 | GGGCCUCGAUCAGAAGGACUUGGGCCCCCCACGA | 28S |
| human-o5 | UCGCUCCCUCGGCCCCGGGAUUCGGCGAGUGC | 28S |
| human-o6 | CCGGGCGCUUGGCGCCAGAAGCGAGAGCCCCUCGGG | 28S |
| human-o7 | GACCGGCUAUCCGAGGCCAACCGAGGCUCCGCGGCG | 28S |
| human-o8 | AGCGACGCUCAGACAGGCGUAGCCCCGGGAGGA | 5.8S |

**Supplementary Table 1:** Oligo sequences (5′ –> 3′) for human oligos (SET-0).

| Oligo ID | sequence (5′ –> 3′) | target rRNA | target pos. |
|---|---|---|---|
| S1_opt-o1 | UGAUCUGAUAAAUGCACGCAUCCCCCC | 18S | 205-231 |
| S1_opt-o2 | CGUGCGAUCGGCCCGAGGUUAUCUAGAGUCACCAA | 18S | 287-321 |
| S1_opt-o3 | AUUCCAUUAUUCCUAGCUGCGGUAUCCAGGCGGCUC | 18S | 837-872 |
| S1_opt-o4 | GGGCCUCGAUCAGAAGGACUUGGGCCCCCCACGA | 28S | 184-217 |
| S1_opt-o5 | UGGCUUCCUCGGCCCCGGGAUUCGGCGAAAGC | 28S | 830-861 |
| S1_opt-o6 | ACGGACGCUUGGCGCCAGAAGCGAGAGCCCCUCGGG | 28S | 3751-3786 |
| S1_opt-o7 | ACCCGGCUAUCCGGGGCCAACCGAGGCUCCUUCGGCG | 28S | 4388-4424 |
| S1_opt-o8 | ACCGACGCUCAGACAGGCGUAGCCCCGGGAGGA | 5.8S | 123-155 |
| **new_S1_o9** | GGCGGACGGGGGGAGAGGGAGAGCGC | 28S | 875-900 |
| **new_S1_o10** | GGCGAGACGGGCCGGUGGUGCGCCCUCGGC | 28S | 1261-1290 |
| **new_S1_o11** | CCAGAAGCAGGUCGUCUACGAAUGGUUUAG | 28S | 4611-4640 |
| **new_S1_o12** | AUCCCCGAUCCCCAUCACGAAUGGGGUUCA | 18S | 1586-1615 |

**Supplementary Table 2:** Oligo sequences (5′ –> 3′) for oligos in SET-1.

| Oligo ID | sequence (5′ –> 3′) | target rRNA | target pos. |
|---|---|---|---|
| S1_opt-o1 | UGAUCUGAUAAAUGCACGCAUCCCCCC | 18S | 205-231 |
| S1_opt-o2 | CGUGCGAUCGGCCCGAGGUUAUCUAGAGUCACCAA | 18S | 287-321 |
| S1_opt-o3 | AUUCCAUUAUUCCUAGCUGCGGUAUCCAGGCGGCUC | 18S | 837-872 |
| S1_opt-o4 | GGGCCUCGAUCAGAAGGACUUGGGCCCCCCACGA | 28S | 184-217 |
| S1_opt-o5 | UGGCUUCCUCGGCCCCGGGAUUCGGCGAAAGC | 28S | 830-861 |
| S1_opt-o6 | ACGGACGCUUGGCGCCAGAAGCGAGAGCCCCUCGGG | 28S | 3751-3786 |
| S1_opt-o7 | ACCCGGCUAUCCGGGGCCAACCGAGGCUCCUUCGGCG | 28S | 4388-4424 |
| S1_opt-o8 | ACCGACGCUCAGACAGGCGUAGCCCCGGGAGGA | 5.8S | 123-155 |
| S1_o9 | GGCGGACGGGGGGAGAGGGAGAGCGC | 28S | 875-900 |
| S1_o10 | GGCGAGACGGGCCGGUGGUGCGCCCUCGGC | 28S | 1261-1290 |
| S1_o11 | CCAGAAGCAGGUCGUCUACGAAUGGUUUAG | 28S | 4611-4640 |
| S1_o12 | AUCCCCGAUCCCCAUCACGAAUGGGGUUCA | 18S | 1586-1615 |
| **new_S2_o11423** | GCUCUGCUACGUACGAAACCCCGACC | 28S | 4640-4665 |
| **new_S2_o25014** | GUGUCGAGGGCUGACUUUCAAUAGAUCG | 28S | 4679-4706 |
| **new_S2_o12152** | AGAUCCAACUACGAGCUUUUUAACUG | 18S | 664-689 |
| **new_S2_o6146** | AACGCCACUUGUCCCUCUAAGAAGU | 18S | 1439-1463 |
| **new_S2_o13419** | CAAGUGCGUUCGAAGUGUCGAUGAUC | 5-8S | 86-111 |

**Supplementary Table 3:** Oligo sequences (5′ –> 3′) for oligos in SET-2.

| Oligo ID | sequence ($5' -> 3'$) | target rRNA | target pos. |
|---|---|---|---|
| S1_opt-o1 | UGAUCUGAUAAAUGCACGCAUCCCCCC | 18S | 205-231 |
| S1_opt-o2 | CGUGCGAUCGGCCCGAGGUUAUCUAGAGUCACCAA | 18S | 287-321 |
| S1_opt-o3 | AUUCCAUUAUUCCUAGCUGCGGUAUCCAGGCGGCUC | 18S | 837-872 |
| S1_opt-o4 | GGGCCUCGAUCAGAAGGACUUGGGCCCCCCACGA | 28S | 184-217 |
| S1_opt-o5 | UGGCUUCCUCGGCCCCGGGAUUCGGCGAAAGC | 28S | 830-861 |
| S1_opt-o6 | ACGGACGCUUGGCGCCAGAAGCGAGAGCCCCUCGGG | 28S | 3751-3786 |
| S1_opt-o7 | ACCCGGCUAUCCGGGGCCAACCGAGGCUCCUUCGGCG | 28S | 4388-4424 |
| S1_opt-o8 | ACCGACGCUCAGACAGGCGUAGCCCCGGGAGGA | 5.8S | 123-155 |
| S1_o9 | GGCGGACGGGGGGAGAGGGAGAGCGC | 28S | 875-900 |
| S1_o10 | GGCGAGACGGGCCGGUGGUGCGCCCUCGGC | 28S | 1261-1290 |
| S1_o11 | CCAGAAGCAGGUCGUCUACGAAUGGUUUAG | 28S | 4611-4640 |
| S1_o12 | AUCCCCGAUCCCCAUCACGAAUGGGGUUCA | 18S | 1586-1615 |
| S2_o11423 | GCUCUGCUACGUACGAAACCCCGACC | 28S | 4640-4665 |
| S2_o25014 | GUGUCGAGGGCUGACUUUCAAUAGAUCG | 28S | 4679-4706 |
| S2_o12152 | AGAUCCAACUACGAGCUUUUUAACUG | 18S | 664-689 |
| S2_o6146 | AACGCCACUUGUCCCUCUAAGAAGU | 18S | 1439-1463 |
| S2_o13419 | CAAGUGCGUUCGAAGUGUCGAUGAUC | 5-8S | 86-111 |
| **new_S3_o6924** | GUCUUCCGUACGCCACAUUUCCCACG | 28S | 141-166 |
| **new_S3_o35101** | GACUGGAGAGGCCUCGGGAUCCCACCUCGG | 28S | 1230-1259 |
| **new_S3_o10474** | ACGAUGAGAGUAGUGGUAUUUCACCG | 28S | 3691-3716 |
| **new_S3_o25177** | UCUAGAAUUACCACAGUUAUCCAAGUAG | 18S | 139-166 |
| **new_S3_o5218** | CUGUAUUGUUAUUUUUCGUCACUAC | 18S | 511-535 |

**Supplementary Table 4:** Oligo sequences ($5' -> 3'$) for oligos in SET-3.

| type | sequence ($5' -> 3'$) |
|---|---|
| 19 nt size selection marker | AGUGUACUCCGAAGAGGAC |
| 32 nt size selection marker | GGCAUUAACGCGAACUCGGCCUACAAUAGUGA |
| $5'$ adapter | GUUCAGAGUUCUACAGUCCGACGAUC |
| $3'$ adapter | 5rApp/UGGAAUUCUCGGGUGCCAAGG/3ddC |
| RTP | GCCTTGGCACCCGAGAATTCCA |
| RP1 forward | AATGATACGGCGACCACCGAGATCTACACGTTCAGAGTTCTACAGTCCGA |
| RPI1 reverse | CAAGCAGAAGACGGCATACGAGATCGTGATGTGACTGGAGTTCCTTGGCACCCGAGAATTCCA |
| RPI2 reverse | CAAGCAGAAGACGGCATACGAGATACATCGGTGACTGGAGTTCCTTGGCACCCGAGAATTCCA |
| RPI3 reverse | CAAGCAGAAGACGGCATACGAGATGCCTAAGTGACTGGAGTTCCTTGGCACCCGAGAATTCCA |
| RPI4 reverse | CAAGCAGAAGACGGCATACGAGATTGGTCAGTGACTGGAGTTCCTTGGCACCCGAGAATTCCA |
| RPI5 reverse | CAAGCAGAAGACGGCATACGAGATCACTGTGTGACTGGAGTTCCTTGGCACCCGAGAATTCCA |
| RPI6 reverse | CAAGCAGAAGACGGCATACGAGATATTGGCGTGACTGGAGTTCCTTGGCACCCGAGAATTCCA |
| RPI7 reverse | CAAGCAGAAGACGGCATACGAGATGATCTGGTGACTGGAGTTCCTTGGCACCCGAGAATTCCA |
| RPI8 reverse | CAAGCAGAAGACGGCATACGAGATTCAAGTGTGACTGGAGTTCCTTGGCACCCGAGAATTCCA |

**Supplementary Table 5:** Size selection marker, adapter and primer sequences ($5' -> 3'$) for experimental methods.
